# Regulation of the one carbon folate cycle as a shared metabolic signature of longevity

**DOI:** 10.1101/2020.07.07.191544

**Authors:** Andrea Annibal, Rebecca George Tharyan, Maribel Fides Schonewolff, Hannah Tam, Christian Latza, Markus Max Karl Auler, Adam Antebi

**Affiliations:** Max Planck Institute for Biology of Ageing, Joseph Stelzmann Str. 9b, 50931, Cologne, Germany; Cologne Excellence Cluster on Cellular Stress Responses in Aging-Associated Diseases (CECAD), University of Cologne, 50931, Cologne, Germany

## Abstract

The metabolome represents a complex network of biological events that reflects the physiologic state of the organism in heath and disease. Additionally, specific metabolites and metabolic signaling pathways have been shown to modulate animal ageing, but whether there are convergent mechanisms uniting these processes remains elusive. Here, we used high resolution mass spectrometry to obtain the metabolomic profiles of canonical longevity pathways in *C. elegans* and identify metabolites regulating life span. By leveraging the metabolomic profiles across pathways, we found that one carbon metabolism and the folate cycle were pervasively regulated in common. We observed similar changes in long lived mouse models of reduced insulin/IGF signaling. Genetic manipulation of pathway enzymes and supplementation with one carbon metabolites reveal that regulation of the folate cycle represents a shared causal mechanism of longevity and proteoprotection.

## Introduction

Cellular metabolism encompasses a highly integrated, complex network that supports the development, growth and reproduction of the organism. Small molecule metabolites comprise basic building blocks for macromolecules and serve as essential carriers of energy and redox potential. Metabolites can also work as signaling molecules that regulate metabolic flux, epigenetic landscapes, gene regulatory networks as well as nutrient and growth signaling pathways^1^. Metabolic dysregulation contributes significantly to diseases such as cancer, cardiovascular disease, and inflammation, and manipulating metabolite levels *in vivo* can help restore metabolic balance and health ^2^.

More recently endogenous metabolites have also emerged as crucial modulators of animal longevity. These include various amino acids, alpha ketoglutarate, spermidine, hexosamines, bile acids, nicotinamides, cannabinoids, ascarosides and other natural compounds that regulate diverse aspects of signaling, metabolism, and homeostasis^3^. Moreover, many of the major longevity pathways are conserved regulators of metabolism, nutrient sensing, and growth ^4^. These pathways include reduced insulin/IGF and mTOR signaling, reduced mitochondrial respiration, dietary restriction mediated longevity and signals from the reproductive system, which remodel metabolism, proteostasis, and immunity towards extended survival ^5^. The question arises, do these diverse signaling pathways converge on shared metabolic outputs that are causal for longevity?

## Results

### Metabolomic fingerprint of long lived mutants

To address this question, we performed high resolution mass spectrometry on several long-lived mutant strains in *C. elegans*, to retrieve unique and common metabolic fingerprints. We used four canonical longevity pathways, namely: insulin/IGF signaling (IIS) deficient *daf-2(e1370)*, dietary restriction (DR) model *eat-2(ad465)*, mitochondrial respiration deficient *isp-1(qm150)* and germline-less *glp-1(e2141)ts* worms. Differentially regulated metabolites were characterized by mass spectrometry-based untargeted metabolomics, using reverse phase liquid chromatography combined with electrospray ionization high-resolution accurate mass (ESI-HRAM) spectrometry. Using this method, we identified and quantified 144 unique metabolites representing different metabolic modules (Fig. 1a, Extended Data Fig. 1a, Supplementary Table 1). PCA analysis of all replicates revealed obvious clustering according to genotype (Extended Data Fig. 1b).

**Figure 1:**
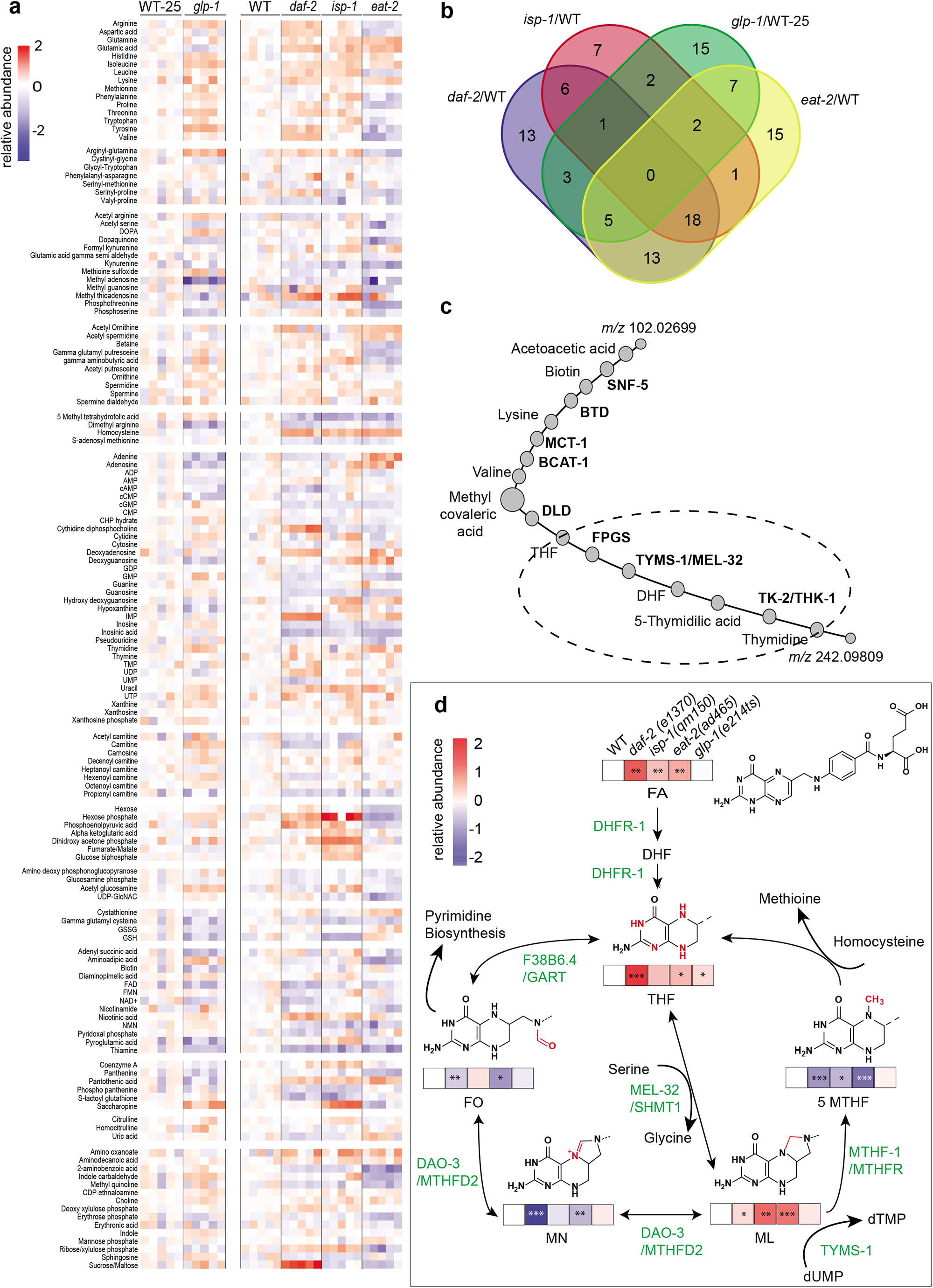
Regulation of the folic acid cycle is altered in longevity mutants. **a**, Untargeted metabolomics analysis of *daf-2(e1370), eat-2(ad465), isp-1(qm150)* and *glp-1(e2141)ts* worms at day 1 of adulthood. *glp-1(e2141)ts* worms are compared with wild type (WT) control, which undergoes the same thermal shift as the mutant. Metabolites concentrations are listed in Supplementary Table 3. Metabolites are manually grouped in different categories. **b**, Venn diagram of the significantly changed metabolites (adj p<.05) for each genotype showing unique and overlapping compounds. **c**, Metabolic-protein network of unknown and known features created by the PIUMet algorithm. Dotted circle indicates the pathway chosen for further investigation. Abbreviation in the chart: SNF-5 (Sodium: Neurotransmitter symporter Family, orthologous to SLC6A8), BTD (biotinidase), MCT-1 (Mono carboxylate Transporter family, an ortholog of human of SLC16A14), BCAT-1 (branched amino acids transporter), DLD (dihydrolipoamide dehydrogenase), FPGS (folylpolyglutamate synthase), TYMS-1 (thymidylate synthetase), MEL-32 (orthologue of SHMT1 serine hydroxymethyl transferase 1) THK-1 (thymidine kinase-1). **d**, Quantification of folic acid intermediates using targeted metabolomic analysis in longevity mutants (day 1). Folic acid, THF and ML accumulate in three out of four longevity mutants. 5MTHF significantly decrease in three longevity mutants. Abbreviation in the chart: FA (folic acid), DHF(dihydrofolic acid), THF(tetrahydrofolic acid), 5MTHF(5-methyl-tetrahydrofolic acid), ML(5, 10-methylene-tetrahydrofolic acid), MN(5, 10-methenyl-tetrahydrofolic acid), FO (formyl-tetrahydrofolic acid). **a,d** N=5 independent biological replicates. **a,d** Metabolite concentrations are normalized to internal standard and converted to log2 for the heat map generation. **a**, Statistics were determined using Fisher test and Benjamini-Hochberg correction (adj p<.05) and **d**, Using one-way-anova and Dunnett’s multiple comparison * p<0.5, ** p<0.01, ***p<0.001 (Supplementary Tables 1 and 3 for statistics).

We first examined the metabolic signatures of each genotype individually. As expected, the relative levels of numerous metabolites were significantly changed (adj p<.05) in *daf-2* (59), *isp-1* (37), *eat-2*(61) and *glp-1*(35), compared to wildtype (WT) (Supplementary Table 1). Some of these changes have been observed in previous studies ^6–8^, confirming the validity of the approach, while others appeared novel (Extended data Fig. 2, Supplementary Table 1). The number of metabolites that were regulated in only one of the genotypes relative to WT was 15 in *glp-1*, 13 in *daf-2*, 15 in *eat-2* and 7 in *isp-1* (adj p<.05) (Fig. 1b, Supplementary Table 1).

Multiple individual metabolites were changed in the each of the pathways (Fig. 1a). Metabolic KEGG pathway enrichment analysis revealed that generally amino acid and nucleotide metabolism were enriched in multiple genotypes (Extended data Fig. 1c-f). Major enriched terms in *glp-1* vs WT-25 comparison included phenylalanine, tyrosine and aspartate metabolism and phosphatidylethanolamine metabolism. *daf-2* vs WT comparisons revealed enrichment in purine, aspartate and alanine metabolism and the malate-aspartate shuffle. *eat-2* vs WT comparisons showed enrichment for glycine and serine metabolism, glutamate, betaine and purine metabolism, and urea cycle. *isp-1* vs WT comparisons showed enrichment in alanine, betaine, tryptophan and glutamate metabolism and oxidation of branched chain fatty acids (Extended data Fig 1c-f).

### Metabolites changed in common across pathways

We next leveraged changes in metabolites across all four pathways to determine if there were any common features (adj p<.05) (Fig. 1b). Although several metabolites showed trends in common across pathways, none emerged as significant from this analysis (Supplementary Table 1). Uracil, isoleucine, glutamine, lysine, appeared higher in all genotypes (isoleucine), but reached significance only in 2 or 3 backgrounds. Erythronic acid, phosphothreonine, propionyl carnitine, kynurenine, gamma-glutamylcysteine, methyl quinoline, glucosamine phosphate, panthenine monophosphate, FAD, thiamine, 2-aminobenzoic acid, S-adenosyl methionine, dimethyl arginine, appeared lower in all genotypes but significant only in 1-3 backgrounds (Extended data Fig. 2).

Because *glp-1* mutants lack germline and often showed dissimilar metabolic features, we limited our search to the metabolites that were commonly regulated (adj p<.05) in three genotypes *daf-2, eat-2 and isp-1*. (Extended data Fig. 2). From this comparison, significantly upregulated metabolites included several amino acids (isoleucine, glutamine, leucine and glutamic acid). Methionine and folate metabolism intermediates were downregulated (5 methyl tetrahydrofolate, S-adenosyl methionine, dimethyl arginine), whereas homocysteine was increased in the three mutants. Nucleotides and related metabolites were also variously dysregulated, (uracil, guanosine, inosine, cyclic GMP, NMN and FAD). Moreover we found changes in phosphothreonine, propionyl carnitine, kynurenine, panthenine monosphoshate, pantothenic acid, glucosamine phosphate, gamma glutamyl cysteine, thiamine and 2-aminobenzoic acid (Extended data Fig 2).

Taking advantage of our unbiased metabolomics acquisition, we additionally retrieved 60 unassigned *m/z* values that were both differentially regulated in all genotypes, (Supplementary Table 3). These uncharacterized *m/z* values were submitted to a pathway predictor software (PIUMet), which uses a machine-learning approach against a database of protein-protein and protein-metabolite interactions to infer a network of dysregulated metabolic pathways.^9^ The network created by PIUMet revealed enrichment in organic acids, branch chain amino acid and folic acid metabolism. Notably, multiple metabolites of the folic acid cycle and associated enzymes were highly enriched (Fig. 1c). Given the biological importance of the folic acid pathway in human health and the large degree of differential regulation we observed in our data (Fig. 1a, Fig. 1c), we decided to follow up on the possible role of these metabolites in longevity.

### One carbon metabolism folate cycle is altered in longevity mutants

One-carbon metabolism mediated by folate cofactors, supports multiple physiological processes including amino acid homeostasis (methionine, glycine and serine), biosynthesis of nucleotides (purines, thymidine), epigenetic maintenance, and redox defense^10–12^. Folate is obtained from the diet and is converted to tetrahydrofolate (THF), which serves as the backbone for one carbon reactions. Enzymes of the folate cycle catalyze the various reactions that transition carbon through three different oxidation states typified by 5,10-methenyl-tetrahydrofolic acid (MN), 5,10-methylene-tetrahydrofolic acid (ML), and 10-formyl-tetrahydrofolic acid (FO) ^13,14^ (Fig. 1d). Many such oxidation and reduction steps are NADPH/NADP dependent, and the folate cycle is actually a major generator of cellular NADPH.

Dihydrofolate reductase (DHFR-1) carries out the first two reaction steps reducing folic acid to dihydrofolic acid (DHF) and on to tetrahydrofolate (THF). Using serine as a methyl donor, serine hydroxy methyl transferase 1 (MEL-32) then produces 5,10-methylene-tetrahydrofolic acid (ML), a key hub intermediate. Thereafter methylene tetrahydrofolate reductase (MTHFR-1) converts ML to 5MTHF, feeding into the methionine cycle, (and reconstituting THF in the process). Methylene tetrahydrofolate dehydrogenase (DAO-3) converts ML to MN, and on to 10-formyl-tetrahydrofolic acid (FO)^13^. Phosphoribosylglycinamide formyltransferase ortholog F38B6.4 transfers the formyl group from FO for use in purine biosynthesis, (and converts FO back to THF in the process). Thymidylate synthase TYMS-1 also uses ML to convert dUMP to dTMP for DNA synthesis, (and in the process restores DHF). DHFR also acts closely with TYMS-1 to recycle DHF back to THF. Notably, DHFR1 and TYMS1 are intimately linked as bifunctional enzymes in parasites, and copurify in plants, and together represent the rate-limiting enzymes for one carbon metabolism ^15–17^

Because our untargeted metabolomic analysis revealed a significant decrease of 5-methyl tetrahydrofolate (5MTHF) in three out of four longevity mutants, we subsequently performed targeted metabolomics and quantified all major folic acid forms, except those conjugated to glutamate. We observed that FA, THF, and ML were more abundant in *daf-2, eat-2* and *isp-1* by 2 to 4-fold. 5MTHF and MN were 2-fold lower in these three genetic backgrounds, while FO was decreased in 3 of 4 genotypes (Fig. 1d). Additionally, THF was significantly elevated in *glp-1* mutants. These results show that metabolites of the FA pathway undergo extensive quantitative changes in multiple long lived strains, and that 5MTHF in particular is robustly and reproducibly reduced.

### Folate cycle enzymes DHFR-1 and TYMS-1 regulate life span

As we observed a common accumulation of folic acid in several long lived worm mutants, we next asked if dietary supplementation of this compound influenced lifespan. Feeding physiological concentrations of folic acid (10 nM), however, had no effect on WT life span, pharyngeal pumping rate or brood size (Extended Data Fig. 3a-c). Bacteria are the main dietary source of folates for worms, but supplementation of submolar FA concentrations significantly did not significantly affect levels of bacterial folate pathway intermediates either (Extended Data Fig. 3d).

We therefore utilized RNAi against key enzymes of the pathway combined with supplementation experiments to manipulate the intracellular concentration of folic acid intermediates. We focused on folate cycle enzymes and first performed a mini-screen using RNAi against *dhfr-1, mel-32, mthf-1, dao-3, tyms-1*, and *F38B6.4* from adult on(Extended Data Fig. 3e). Only *tyms-1i* and *dhfr-1i* increased mean lifespan significantly, by +25% and +32%, respectively. Further ageing experiments confirmed that *dhfr-1* RNAi administered from adult on consistently extended worm mean life span by 17-22% (Fig. 2a), with little effect on food intake as measure by pharyngeal pumping rates or reproductive capacity as measured by brood size (Extended Data Fig. 3f,g). Similarly, RNAi knock down of *tyms-1* also led to a 20% increase in worm median lifespan (Fig. 2a), consistent with the close association with *dhfr-1*.

**Figure 2:**
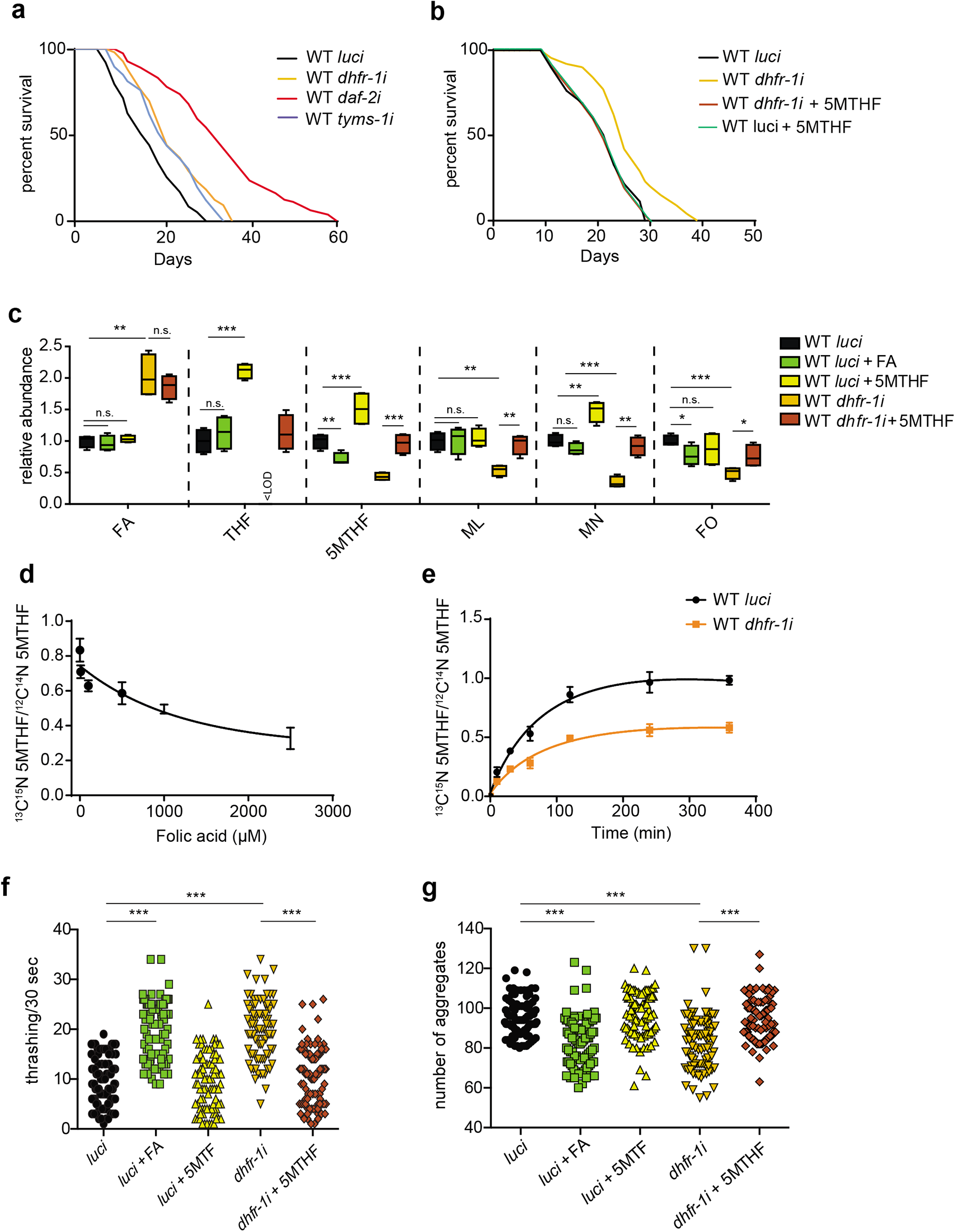
*dhfr-1i* prolongs nematode lifespan in a 5MTHF dependent manner. **a**, *dhfr-1* and *tyms-1* RNAi treatment increase wild type lifespan (from L4 stage). **b**, Supplementation of 10 nM of 5MTHF is sufficient to abolish *dhfr-1i* longevity (from L4 stage). **c**, quantification of folic acid intermediates using targeted mass spectrometry in wild type worms (day 1) with *luci* and *dhfr-1i* treatment in the presence or absence of folic acid and 5MTHF. *dhfr-1i* increases folic acid and decreases 5MTHF and downstream intermediates. **d**, Increasing concentrations of FA inhibit 5MTHF incorporation (day1). **e**, Incorporation of labeled 5MTHF over time in worms with *luci* (black line) and *dhfr-1i* (orange line) (day 1). **f**, Thrashing assay of polyQ35 worms at day 7 of adulthood. **g**, Protein aggregate quantifications in polyQ40 model in worms at day 7 of adulthood. *dhfr-1i* as well as folic acid supplementation have beneficial effect, whereas 5MTHF is detrimental. **a,b**, n=150 per repeat per condition. **c**, N=5 independent biological replicates. **d,e** N=4 independent biological replicates. **f,g** n=30 worms, N=3 independent biological replicates. **a,b**, Statistics were by two-sided Mantel–Cox log-rank test. **c,f,g**, Significance was assessed using one-way anova and Dunnett’s multiple comparisons test. * p<0.5, ** p<0.01, ***p<0.001. (Supplementary Table 3 for statistics)

We then measured the levels of folate cycle intermediates upon *dhfr-1i* knockdown. *dhfr-1* RNAi led to an accumulation of upstream FA, concomitant with a reduction in downstream intermediates THF, 5MTHF, MN, ML, and FO, confirming our previous results^18^. Consistent with a critical role as rate-limiting enzyme in the one-carbon cycle, supplementation of *dhfr-1i* treated worms with the downstream intermediate 5MTHF restored near normal levels of all folate cycle intermediates, except for upstream FA which remained high. In *luciferase (luci)* RNAi controls, dietary 5MTHF also elevated levels of THF, 5MTHF, MN beyond untreated animals, but had no effect on levels of ML or FO (Fig. 2c). Given that 5MTHF uptake by the worm largely restored folate pools, we next asked whether supplementation would impact longevity. Whereas 10 nM 5MTHF supplementation had no effect in *luci* controls, it abolished extension of life by *dhfr-1i* (Fig. 2b), indicating that *dhfr-1i* longevity arises from lower levels of 5MTHF (or altered levels of other folate intermediates) in the worm.

### FA regulates 5MTHF uptake

Interestingly, we noticed that *dhfr-1i* led to accumulation of FA, and that FA supplementation led to similar changes in folate intermediates as *dhfr-1i* knockdown in the worm, though not to the same extent. Specifically, both conditions led to a lowering of 5MTHF, MN and FO (Fig. 2c). This suggested that the FA-induced decrease in these downstream intermediates might arise via inhibition of uptake or negative feedback. To test this idea, we measured the incorporation of 5MTHF in adult worms, by quantifying the ratio of ^13^C and ^15^N double-labeled versus non-labeled 5MTHF. We found that both increasing concentrations of FA as well as *dhfr-1* RNAi counteracted 5MTHF incorporation *in vivo* (Fig. 2d,e). These findings suggest an inhibitory role of FA in the assimilation of folate cycle intermediates, supporting previous *in vitro* work ^19,20^.

### High FA and low 5MTHF ameliorate models of polyQ proteotoxicity

Many long lived strains exhibit improved protein homeostasis that is often manifest as greater resistance to toxic aggregate-prone proteins during ageing. Various polyglutamine repeats expressed in *C. elegans* muscle, modeling Huntington’s disease, form aggregates and induce age-related progressive paralysis, with severity and age of onset related to repeat length^21^. To explore other possible benefits of FA supplementation or *dhfr-1i* treatment, we investigated the effect of folates on polyQ repeat proteotoxicity models. Whereas supplementation of FA or endogenous accumulation of FA via *dhfr-1i* significantly enhanced the motility of polyQ35 worms, supplementation with 5MTHF in both conditions was detrimental (Fig. 2f). We also tested the polyQ40 proteotoxicity model, and found that *dhfr-1i* reduced the number of visible protein aggregates, while 5MTHF supplementation brought such aggregates back to WT levels (Fig. 2g). These findings suggest that elevated FA and lower 5MTHF improve protein quality control and ameliorate proteotoxicity.

### *dhfr-1i* induces methionine restriction

5MTHF provides essential substrates for methionine synthetase (MS) and is a main source of carbon units for the methionine cycle^10^. Homocysteine captures the methyl group from 5MTHF to form methionine and is rapidly converted to S-adenosyl methionine (SAM). SAM then donates its methyl group to various molecular acceptors, and in the process generates adenosyl homocysteine, which is then converted to homocysteine. Methylation of homocysteine completes the methionine cycle (Fig. 3a). Aside from the methionine cycle, homocysteine can be channeled into transsulfuration, glutathione and pyruvate pathways (not shown).

**Figure 3:**
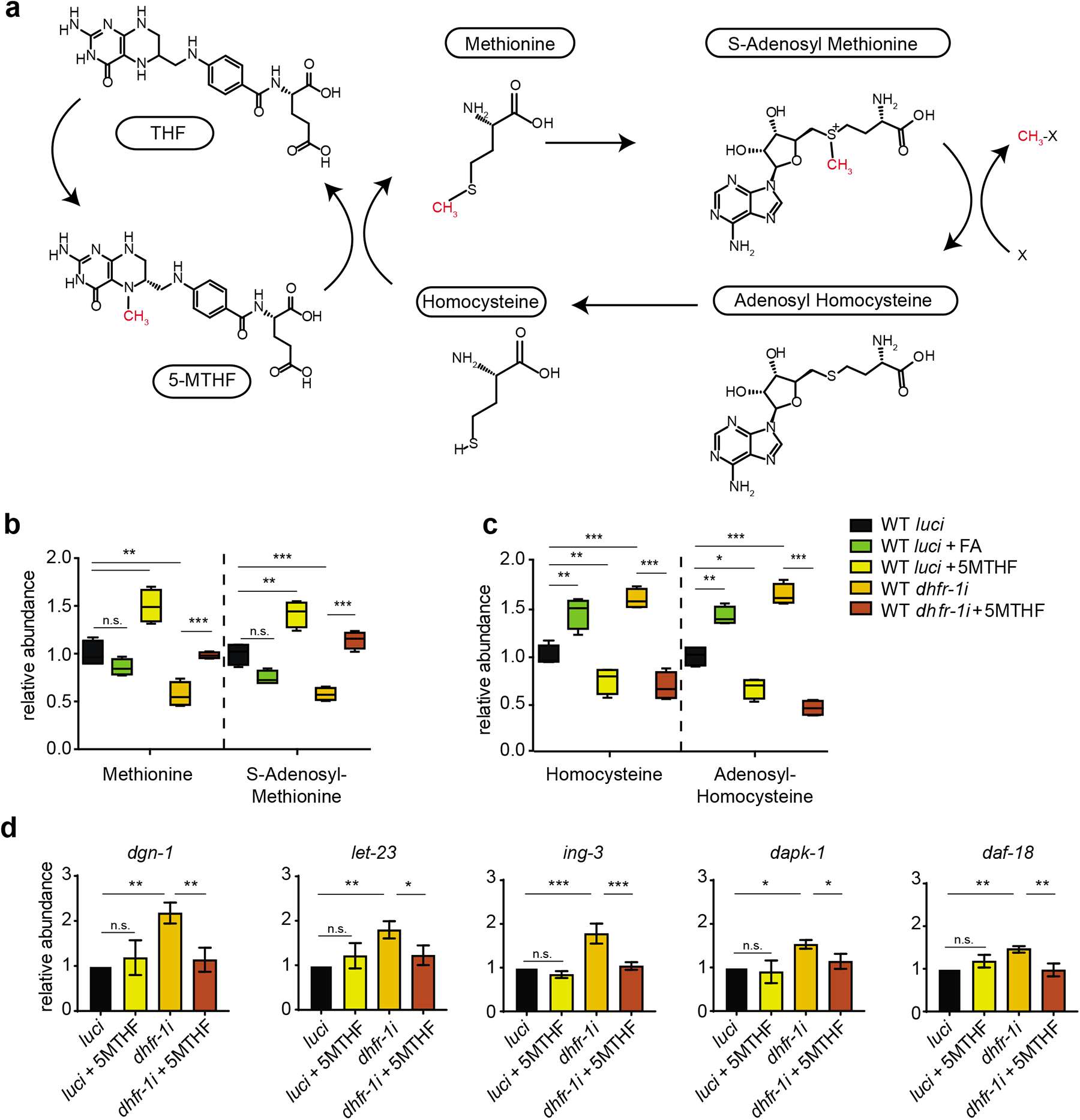
*dhfr-1i* affects methionine cycle and mimics methionine restriction. **a**, Schematic of methionine cycle. **b,c** Quantification of methionine cycle intermediates using targeted mass spectrometry in *dhfr-1i* and *luci*, with or without FA and 5MTHF supplementation (day 1). *dhfr-1i* decreases methionine and S-adenosyl methionine levels and increases homocysteine and adenosyl-homocysteine levels. This increase is reversed by supplementation with 5MTHF. **d**, Relative gene expression of specific gene signature for methionine restriction (day 1). *dhfr-1i* increases the levels of *dgn-1, let-23, ign-3, dapk-1*, and *daf-18*. **b, c**, N=5 independent biological replicates. **d**, N=3 independent biological replicates. **b,c,d** Significance was assessed using one-way anova with Dunnett’s multiple comparisons test.* p<0.5, ** p<0.01, ***p<0.001.

We next investigated how levels of methionine cycle intermediates varied upon *dhfr-1i* and 5MTHF feeding using targeted metabolomics. Notably, *dhfr-1* knockdown resulted in significantly lower levels of methionine and S-adenosyl methionine. *dhfr-1* knockdown or FA supplementation also resulted in accumulation of homocysteine and adenosyl-homocysteine. Supplementation of 10 nM 5MTHF was sufficient to significantly restore methionine levels in *dhfr-1i* (Fig. 3b,c), and reduce levels of homocysteine and S-adenosylhomocysteine (Fig. 3b,c). Thus *dhfr-1i* and 5MTHF supplementation impact methionine pools *in vivo*.

Methionine restriction induces a characteristic transcriptional profile in mammalian cells that can be used to monitor the process^22^. We used a *C. elegans* methionine restriction model, *metr-1i*, to characterize this state in the worm and validated that several nematode orthologues of genes found in the mammalian study were similarly upregulated (Extended data Fig. 3i). To test whether *dhfr-1i* knockdown induces the same signature, we tested the mRNA expression levels of these representative genes and observed a significant 1.5-2-fold increase upon *dhfr-1i*. This increase was specific, as 5MTHF supplementation of *dhfr-1i* treated animals reverted expression to control levels (Fig. 3d). Taken together, these findings reveal that *dhfr-1i* not only reduces 5MTHF, but methionine and SAM as well, thereby activating a specific gene response indicative of methionine restriction ^22,23^.

To investigate more globally how *dhfr-1i* affects other metabolic pathways, we performed untargeted metabolomics on *dhfr-1i* with and without 5MTHF supplementation (Extended Data Fig. 4a, Supplementary Table 2). *dhfr-1i* exhibited specific metabolic signatures associated with methionine restriction, including lower levels of tryptophan and methionine sulfoxide (adj p<.05) (Extended Data Fig. 4b) ^24,25^. In addition, we observed changes in nucleoside and energy metabolism. Levels of thymidine and uracil were significantly lower, whereas AMP and NMN were significantly higher (adj p<.05) upon *dhfr-1i. dfhr-1i* treated animals also showed significantly lower levels of acetyl and propionyl carnitine and increased levels of alpha ketoglutarate and dihydroxy acetone phosphate, possibly indicating changes in lipid metabolism, TCA cycle and glycolysis. Furthermore, spermidine metabolism was also altered by *dhfr-1i* as acetyl putrescine and spermine were increased (Extended Data Fig. 4b)^26^.

Interestingly, many of these changes described above were either partially or fully reversed in the presence of 5MTHF, though not all changes reached significance (Supplementary Table 2). These data reveal possible unexpected connections between the one carbon metabolism and other metabolic processes.

### 5MTHF affects the longevity of long lived IIS and mitochondrial mutants

Our original metabolomic analysis indicated that *C. elegans* longevity mutants, *daf-2* and *isp-1* exhibit a prominent reduction in 5MTHF and other FA intermediates (Fig. 1d). We asked whether lower levels of 5MTHF play a role in life span regulation in these mutants by supplementing 5MTHF. 5MTHF treatment modestly reduced mean and max life span of *daf-2* (mean: −15.8%, max: −22%) and *isp-1* (mean: −13%, max:-15%), suggesting that lower levels of the 5MTHF contribute towards longevity in these mutants(Fig. 4a,b).

**Figure 4:**
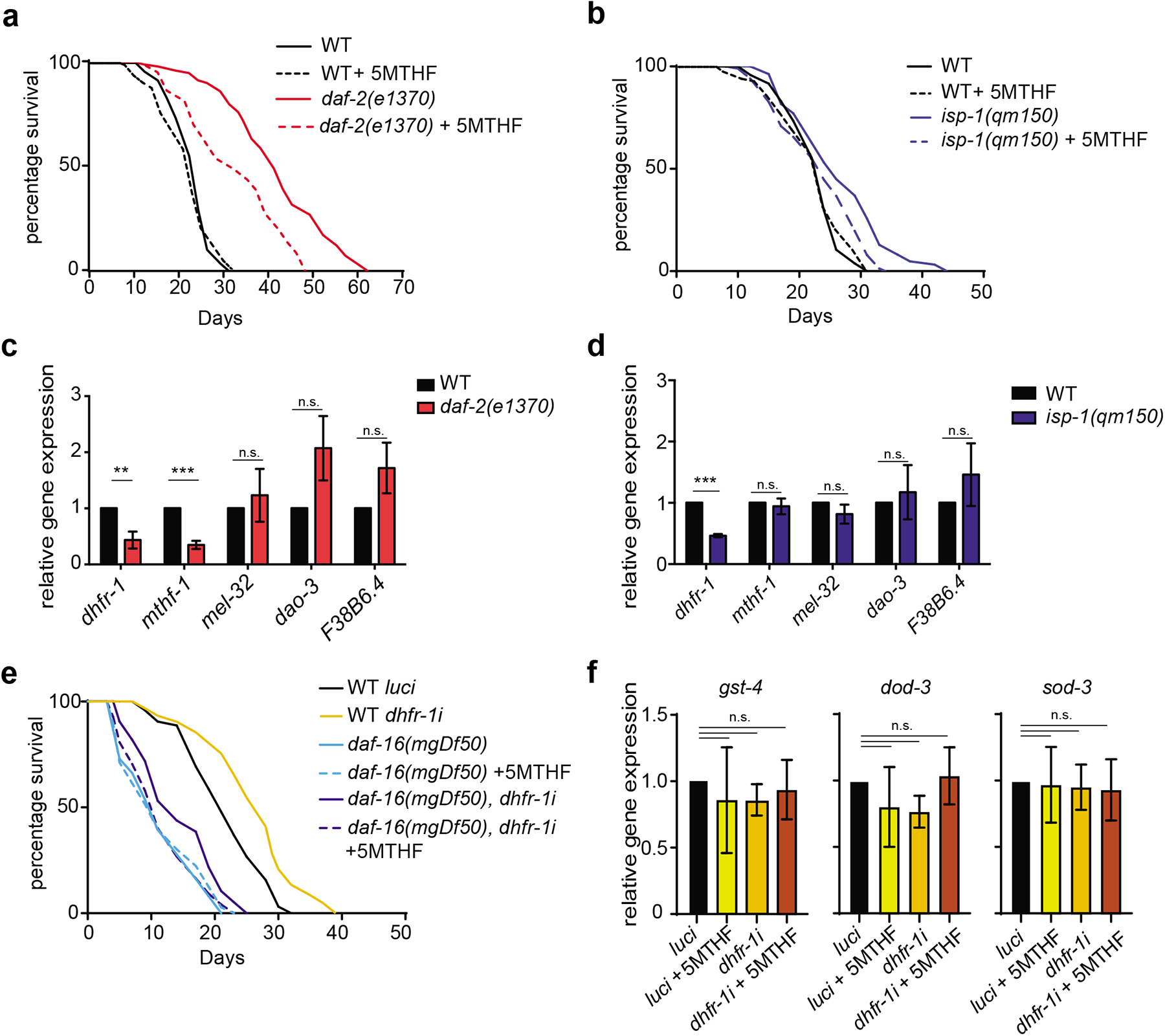
*dhfr-1* and 5MTHF act within the insulin/IGF and mitochondrial signaling pathways. **a**, Lifespan of *daf-2(e1370)* supplemented with 10nM 5MTHF (from L4 stage). **b**, Lifespan of *isp-1(qm150)* upon 5MTHF supplementation (from L4 stage). 5MTHF does not alter WT lifespan but is detrimental to *daf-2* and *isp-1* longevity. **c,d** Folic acid cycle gene expression in *daf-2(e1370)* and *isp-1(qm150)* (day 1 adult). *dhfr-1* expression is lower in both genotypes. **e**, Lifespan experiment of wild type and *daf-16(mgDf50)* with *luci* and *dhfr-1i* in the presence or absence of 5MTHF (from L4 stage). *dhfr-1i* extended *daf-16* mutant lifespan and supplementation with 5MTHF abolished it. **f**, *daf-16* dependent gene expression in wild type worms with *luci* or *dhfr-1i* in the presence or absence of 5MTHF (day 1). No significance (n.s.) difference was found between the conditions. **a,b,e** n=150 per repeat per condition. **c,d,f** N=3 independent biological replicates. **a,b,e**, statistics were by two-sided Mantel–Cox log-rank test. **c,d,f**, Significance was assessed using one-way anova and Dunnett’s multiple comparisons test. * p<0.5, ** p<0.01, ***p<0.001 (Supplementary Table 3 for statistics).

To further investigate the cause of the lower 5MTHF level in these two genotypes, we quantified the mRNA expression of various folic acid cycle genes. Interestingly, *dhfr-1* mRNA expression was decreased by more than 50% in both mutants. *mtfh-1* expression was also ca. 50% lower in the *daf-2* background. (Fig. 4c,d). Other folic acid cycle genes (*mthr-1, mel-32, dao-3* and *F38B6.4*) were unchanged.

The DAF-16/FOXO winged helix transcription factor is a major regulator of longevity whose mutation negates life extension in both *daf-2* and *isp-1* pathways ^27,28^. In response to reduced insulin/IGF or mitochondrial signaling, DAF-16 localize to the nucleus to control transcription of target genes involved in oxidative stress response, heat shock, and lipogenesis^29^. Given that *dhfr-1* is a regulatory target of these pathways, we asked if *dhfr-1i* lifespan extension also shows *daf-16* dependence. To our surprise, ageing experiments showed that the median lifespan of *daf-16(mgDf50)* was modestly increased upon *dhfr-1i* by 6-8% (Fig. 4e). This increase was reversed upon supplementation with 5MTHF. These findings suggest that *dhfr-1i* lifespan extension may act downstream or parallel to *daf-16*. To further explore this idea, we monitored the expression of specific *daf-16* dependent genes, *gst-4, dod-3* and *sod-3*^30^. Their expression was unaltered upon *dhfr-1i* and 5MTHF treatment, reinforcing the notion that the *dhfr-1i* lifespan extension is downstream or parallel to *daf-16* (Fig. 4f).

We also asked whether *dhfr-1i* affects *daf-2* longevity (Extended Data Fig. 3h). However, ageing experiments showed no difference in mean and max lifespan in the *daf-2* background upon *dhfr-1i*. Altogether these findings suggest that *dhfr-1* is an output of the insulin/IGF pathway, acting downstream or parallel to *daf-16*.

### Mammalian insulin pathway regulates folic acid cycle intermediates

The insulin receptor substrate protein 1 (*Irs1*), is a key mediator of insulin/IGF signaling (IIS) in mammals. Homozygous disruption of *Irs1* in mice results in lifespan extension, resistance to several age-related pathologies including bone and motor dysfunction, skin, and protection against glucose intolerance^31,32^. Because reduced insulin signaling in worms altered folate homeostasis, we wondered if similar changes occur in the *Irs1*^-/-^ mouse model. Targeted metabolomics of folic acid intermediates using brain and liver tissues from male *Irs1*^-/-^ knock out mice revealed a profile of intermediates similar to worm *daf-2* mutants (Fig. 5a,c). Folic acid and THF were increased up to 2-fold in both brain and liver in *Irs1^-/-^*. ML tended to increase as well but did not reach significance. In addition, 5MTHF, MN, and FO were significantly decreased. We also quantified methionine cycle intermediates in these tissues and observed significantly lower levels of methionine and S-adenosyl methionine, and a large increase in homocysteine (Fig. 5b,d). These data suggest that the insulin/IGF signaling pathway similarly regulates the folate and methionine cycle from worms to mammals.

**Figure 5:**
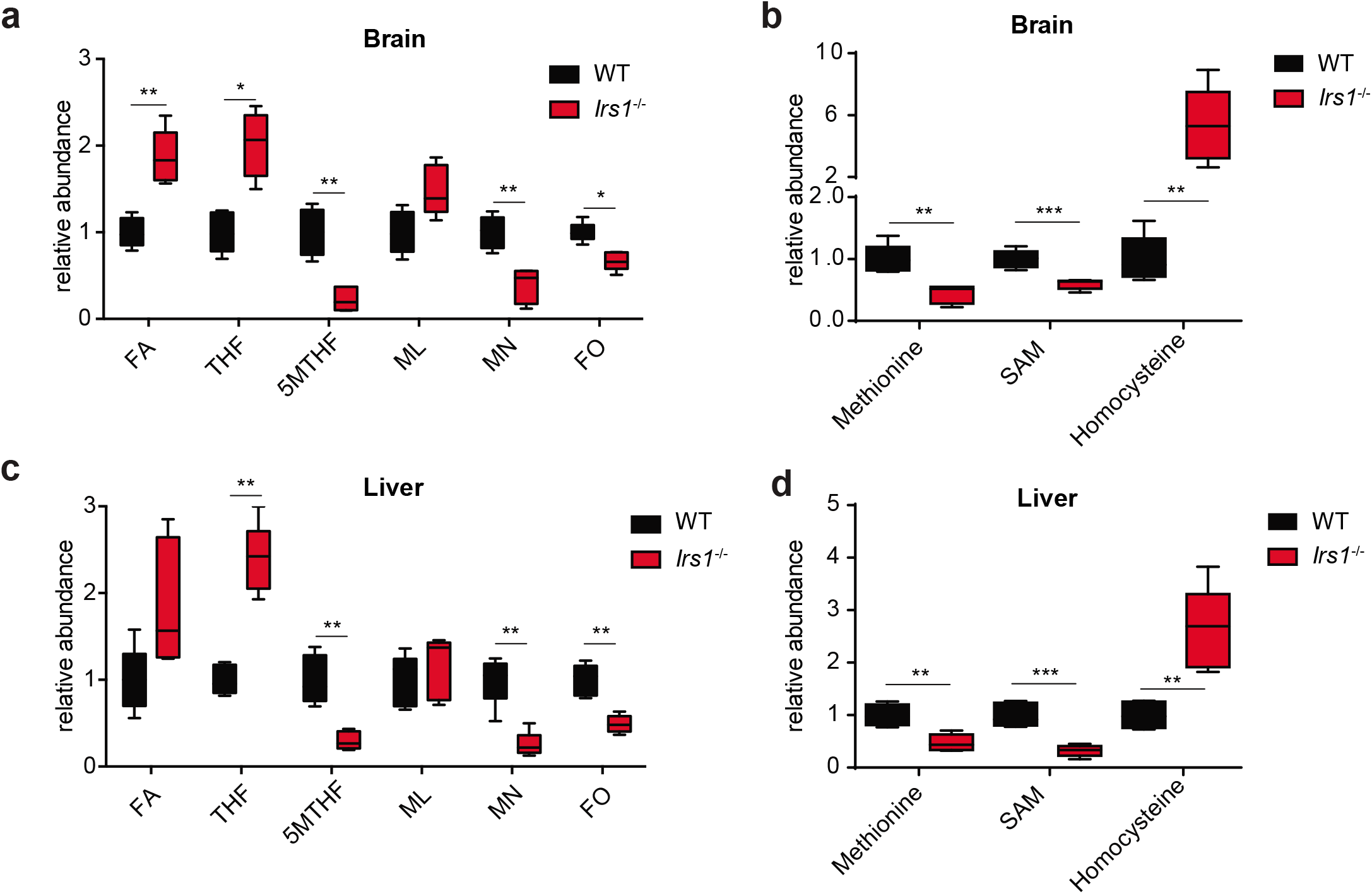
Reduced insulin/IGF signaling alters folic acid intermediates in mice. **a,c**, Targeted metabolomics of folic acid intermediates in brain and liver in wild type and *Irs1^-/-^* full body knock out mouse. Folic acid and THF increased in *Irs1^-/-^*, whereas 5MTHF is lower in the mutant. **b,d**, Quantification of methionine cycle intermediates in brain and liver in wild type and *Irs1^-/-^* full body knock out mouse. Methionine and SAM are lower in the mutants and homocysteine is higher. **a-d**, N=5 mouse, independent biological replicates. **a,d**, Significance was assessed using one-way anova Dunnett’s multiple comparisons test. * p<0.5, ** p<0.01, ***p<0.001 (Supplementary Table 3 for statistics).

## Discussion

Folates are essential vitamins derived from dietary sources and the microbiome. The folate cycle provides one carbon units for an extensive metabolic network that fuels the methionine cycle, transsulfuration pathway, *de novo* purine synthesis, thymidine production, serine, glycine, glutathione and NADPH pools, and thereby regulates cellular redox state, growth, and proliferation^13,33^. The major canonical longevity pathways also profoundly regulate metabolism, growth and organismal physiology in numerous ways^34^. By generating the metabolic profiles of several different long-lived mutants in *C. elegans*, we discovered that folate cycle intermediates and their enzymes represent a convergent focal point for longevity across conserved signaling pathways.

Generally, we found that several folate intermediates were consistently altered. 5-methyltetrahydrofolate levels were specifically decreased in long-lived models of downregulated insulin/IGF signaling (*daf-2*), mitochondrial respiration (*isp-1*), and dietary restriction (*eat-2*), while THF was upregulated in reproductive signaling (*glp-1*) mutants. In accord with a physiological role in longevity, knockdown of folate cycle enzyme *dhfr-1* in the wild type background was sufficient to similarly decrease 5MTHF, extend life span and reduce polyQ proteotoxicity. Supplementation with this metabolite restored the folate pool and reversed these phenotypes. Consistently, *daf-2*/InsR and *isp-1*/Rieske longevity was reduced by 5MTHF supplementation, and these mutant strains downregulated levels of *dhfr-1* mRNA, supporting regulation of folate metabolism by these signaling pathways. Strikingly we saw similar changes in folate cycle intermediates in tissues of Irs1-/- knockout mice and *daf-2/InsR* mutant worms, revealing that the control of the folate cycle by insulin/IGF signaling is evolutionarily conserved. Thus, reduced 5MTHF and other folate intermediates are causal and/or associated with longevity as part of a shared mechanism in several pathways.

Previous studies in *C. elegans* have described disparate effects of FA on longevity. The anti-diabetic drug metformin extends *C. elegans* life span by disrupting microbial folate production and inducing methionine restriction ^35^, while inhibition of endogenous *C. elegans* genes for folate uptake or folate polyglutamase activity had little effect on longevity ^36^. By contrast, Rathor and colleagues reported that uM levels of FA extend life span ^37^. We saw no effect of 10 nM FA supplementation on worm life span, but observed an improvement in polyglutamine proteotoxicity models, suggesting folates as potential therapeutics for proteoprotection. These varied outcomes probably reflect differences in compound dose and availability on uptake or feedback, as well as the complex interplay between diet, microbiome and host ^13,38^. In our case, we targeted rate limiting interlinked enzymes of the *C. elegans* folate pathway, *dhfr-1* and *tyms-1*, and saw coherent changes in folate intermediates and extension of worm life span. Supplementation with nanomolar amounts of 5MTHF were sufficient to restore folate pools of *dhfr-1i*, with little effect on bacterial folates pools. Knockdown of other folate cycle enzymes might not have elicited these phenotypes in our hands because of RNAi efficiency or pleiotropic effects on other processes. Interestingly, a link between longevity and folate metabolism has been recently reported in budding yeast: Rpl22 ribosomal protein mutants increase yeast replicative life span, show lower levels of metabolites associated with folate, serine and methionine metabolism, and deletion of 1C enzymes enhanced wild type longevity ^39^.

Untargeted metabolomic analysis of *dhfr-1i* also revealed changes in a handful of other metabolic pathways, which showed partial reversal upon replenishing 5MTHF. As expected, we saw significant changes in folate intermediates, methionine cycle, and pyrimidine, and polyamine metabolism. Interestingly, we also observed unexpected changes in AMP, cAMP, NMN, as well as propionyl and acetylcarnitines, alpha ketoglutarate and dihydroxyacetone phosphate levels, suggesting an impact on TCA, glycolysis, fat, and energy metabolism. Interestingly, elevated alpha ketoglutarate has been suggested to promote longevity by downregulation of mTOR signaling ^40^. These findings illustrate the power of this approach to illuminate the proximal and distal connectivity of the folate cycle on metabolism.

An important unresolved question is what is the mechanism by which *dhfr-1i* and 5MTHF reduction trigger longevity? Because *dhfr-1* knockdown altered several folate intermediates, longevity could arise from metabolic pathways emanating from multiple branchpoints. Among them, we observed that *dhrf-1* knockdown perturbs the methionine cycle, leading to reduced levels of methionine and S-adenosylmethionine and elevated levels of homocysteine, suggesting methionine restriction as one of several possible mechanisms. Consistently, we observed changes in gene expression associated with this process. Methionine restriction promotes longevity in diverse species, and can lead to lower levels of protein synthesis directly through amino acid limitation, or indirectly through general control and mTOR signaling ^41^. As a potent methyl donor, SAM catalyzes methylation of rRNA, DNA, epigenetic factors, and spermidine synthesis. Homocysteine can also be diverted to transsulfuration reactions, producing cysteine, H_2_S, and glutathione, and itself can modulate insulin/IGF signaling ^42^. These and related processes have been implicated in life span regulation in diverse contexts ^33,43–45^. Indeed, folate cycle and methionine restriction affect diverse aspects of mammalian physiology including immunity, inflammation, cancer, and progeria ^46–49^. Folate supplementation has also been shown to modulate the DNA methylation ageing clock ^11^. Thus, precise manipulation of folate intermediates may provide new entry points to broadly improve human health and disease.

Metabolomics has emerged as a powerful approach to identify not only markers of health, disease, and ageing, ^50^ but also causal mechanisms ^51^. Our analysis of the different longevity pathways revealed congruent changes in other crucial metabolic pathways ^52^. These profiles can serve as a useful resource for the field. Among the changes, we saw modulation of kynurenine and nicotinamide metabolism, which are biochemically interlinked, and have been previously shown to regulate longevity ^53,54^. We detected consistent changes in various nucleoside related metabolites involved in nucleic acid synthesis and signaling, some of which have also been associated with longevity (thymine, adenosine ^55,56^). As well, we observed changes in proprionyl carnitine, which impacts acyl-CoA and succinate metabolism, and gamma glutamylcysteine involved in glutathione metabolism ^57^. We also saw changes in 2-aminobenzoic acid in three longevity mutants. Derived from tryptophan by action of the kynurenine pathway, this metabolite has been suggested as endogenous marker of organismal mortality in the nematode ^58^. It will be interesting to validate these various changes by targeted metabolomics, and determine the potential roles of these and related metabolites in impacting health and life span.

In sum these studies reveal the power of leveraging multiple longevity pathways to uncover molecules that can impact the ageing process and suggest several new hypotheses to be tested. Our discovery that the folate cycle is a convergent mechanism in multiple longevity pathways, and whose regulation by insulin/IGF signaling is conserved in evolution, could provide new ways to improve health during ageing.

## Contributions

A. Antebi and A. Annibal designed experiments and wrote the paper. A. Annibal carried out all experiments. R.G.T. performed *isp-1* supplemented with 5MTHF lifespan and *isp-1* qPCR. M.F.S. performed flux analysis experiments. H.T. performed q35 and q40 experiments and *daf-2(e1370) dhfr-1i* lifespan. M.M.K.A. prepared mouse samples and extracted metabolites from tissues. M.F.S, H.T and C.L. gave technical assistance.

## Ethics declarations

Mouse tissues were kindly provided by Linda Partridge. This study was performed in strict accordance with the recommendations and guidelines of the Federation of European Laboratory Animal Science Associations (FELASA). The protocol was approved by the “Landesamt fuer Natur, Umwelt und Verbraucherschutz Nordrhein-Westfalen”.

## Acknowledgements

We thank Dr. Orsolysa Symmons for critical reading of the manuscript and the MPG for research support. We additionally thank the bioinformatics core facility for assistance (MPI-AGE).

---

**Extended data Fig.1:**
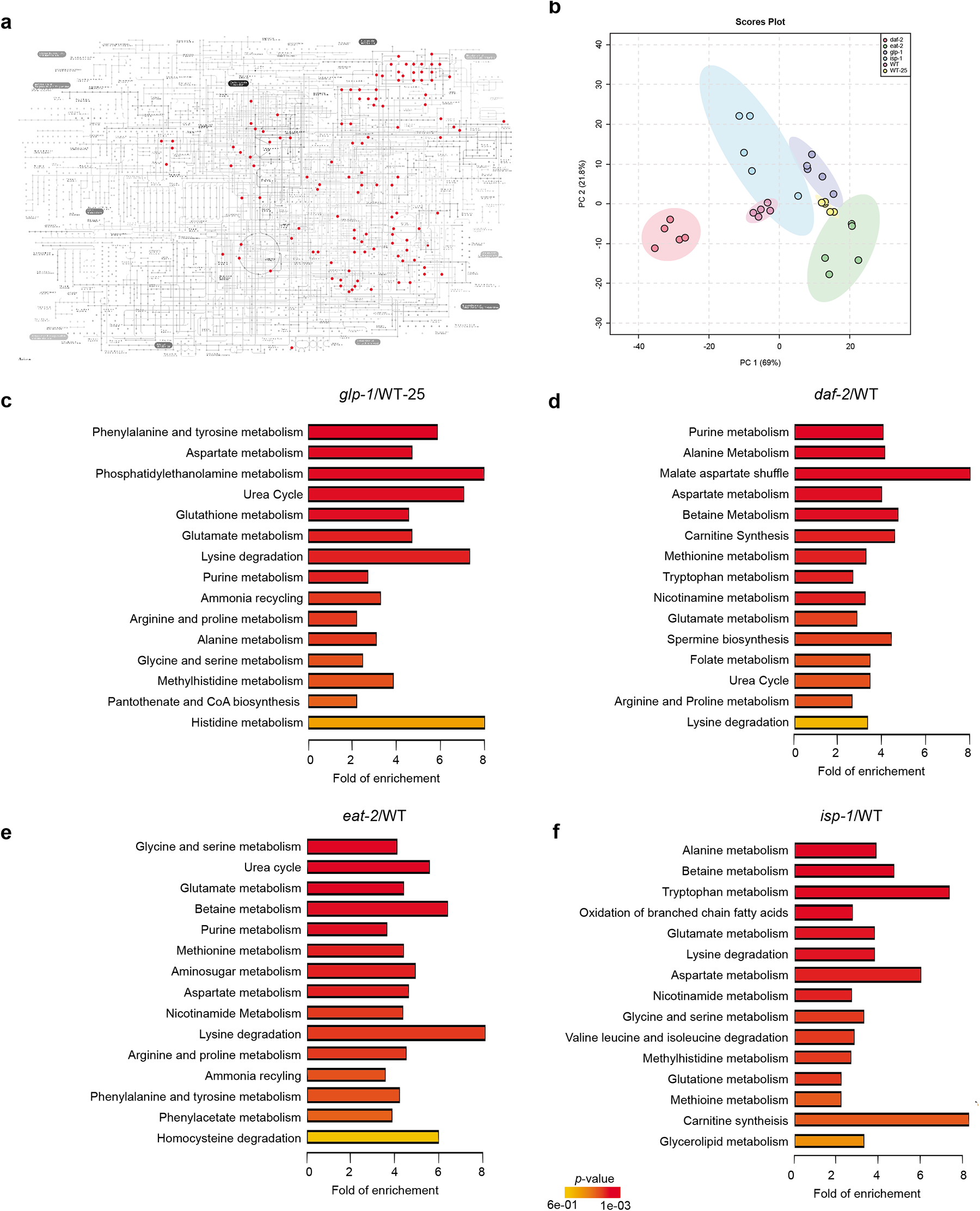
Untargeted metabolomics and pathway analysis of longevity mutants (linked to Fig.1). **a**, Mapping of identified metabolites onto KEGG pathways. **b**, PCA scores of the untargeted metabolomics features of all five conditions obtained by uploading raw files to MetaboAnalyst(https://www.metaboanalyst.ca). **c-f**, Enrichment analysis of significant metabolites (adj p<.05) obtained by uploading the differential regulated metabolites to MetaboAnalyst (https://www.metaboanalyst.ca) in different mutants compared to WT.

**Extended data Fig.2:**
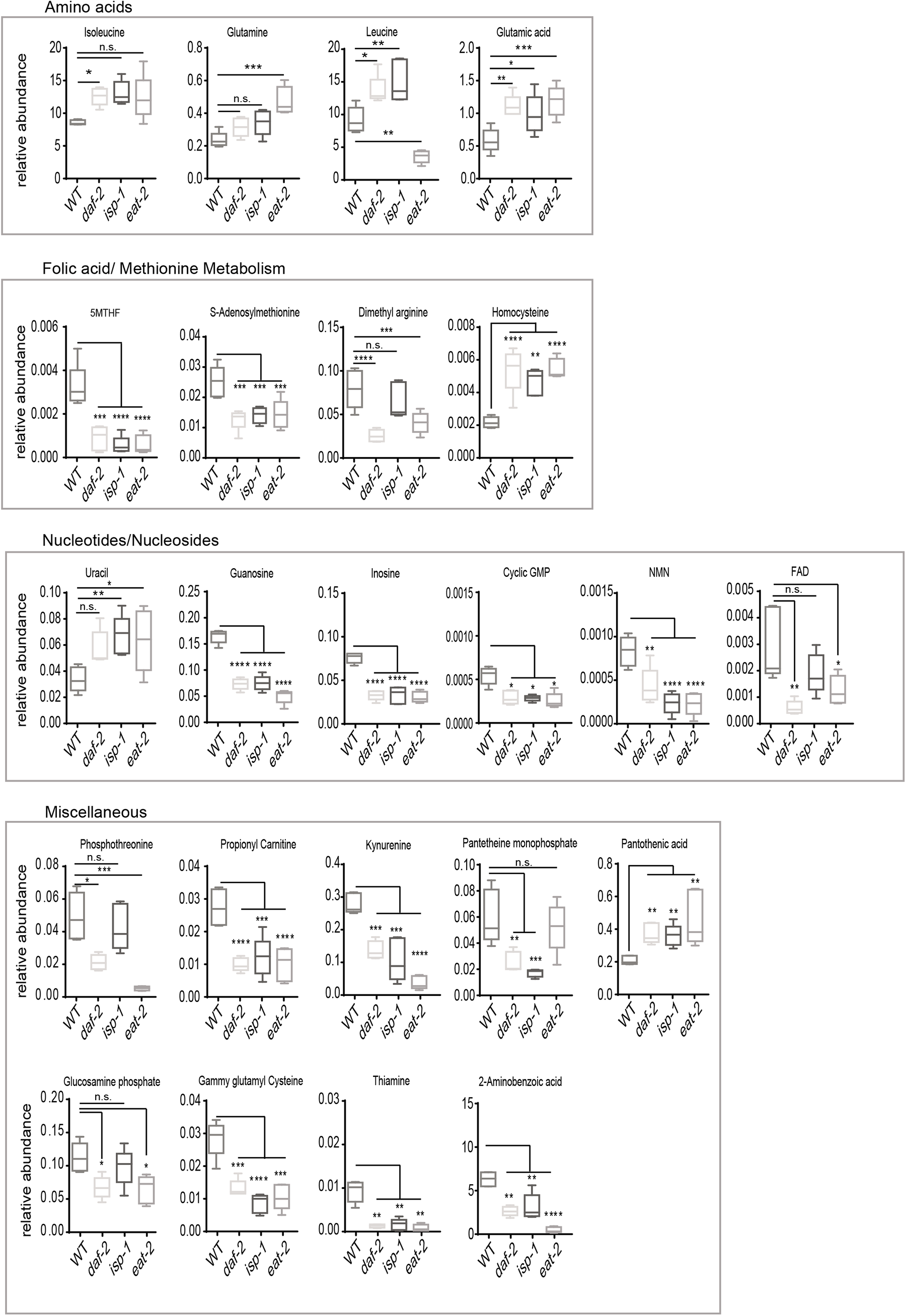
Differentially regulated metabolites in the longevity mutants (linked to Fig. 1). Significant metabolites (adj p<.05) are selected from the untargeted metabolomics analysis (Fig.1a, Supplementary Table 1) and plotted. Significance was assessed using one-way anova Dunnett’s multiple comparisons test * p<0.5, ** p<0.01, ***p<0.001.

**Extended data Figure.3:**
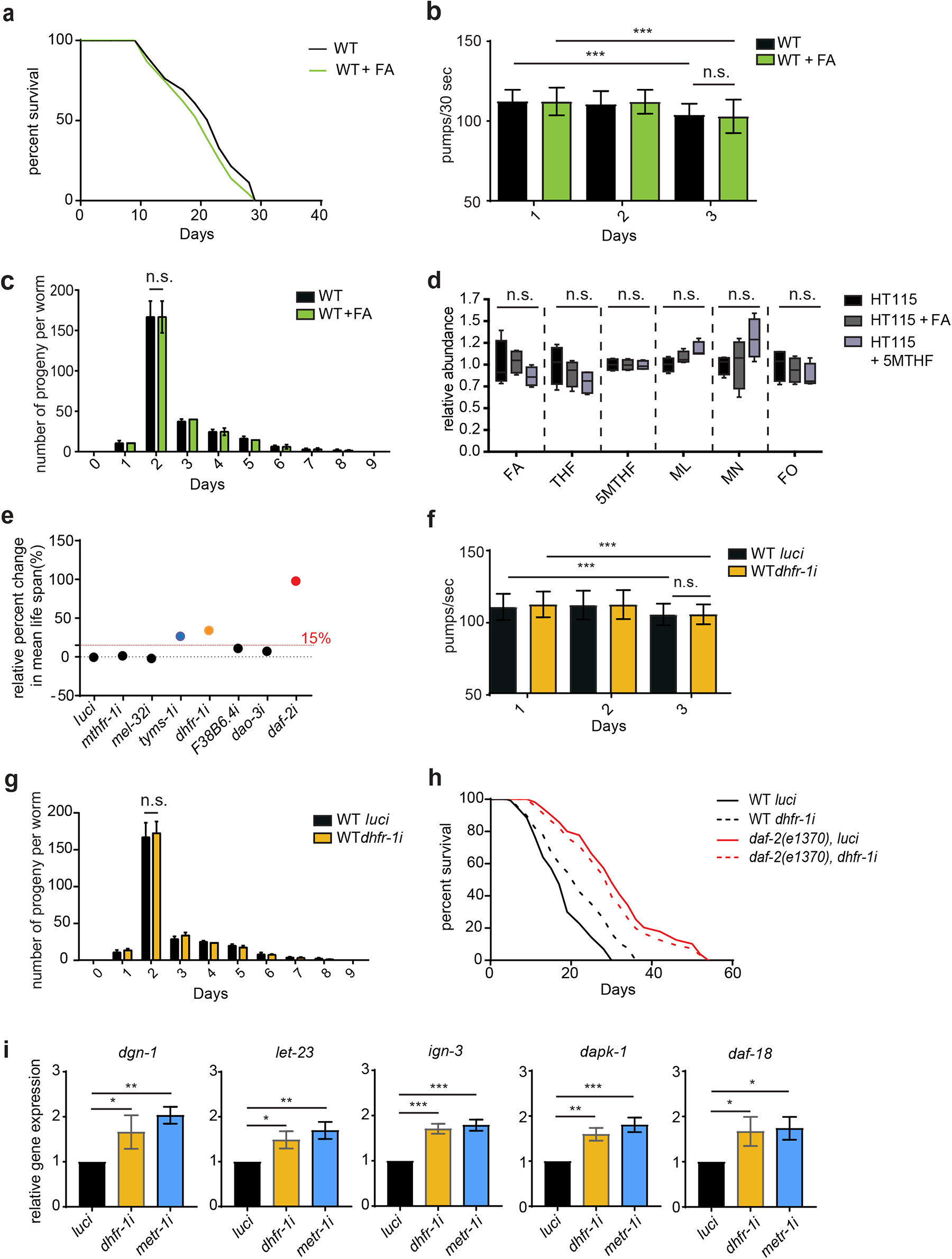
Effect of Folic acid supplementation on worms and investigation on the role of folic acid cycle gene in lifespan (linked to Fig. 2,3,4). **a**, Supplementation of 10 nM of folic acid does not have any effect on WT lifespan (from L4 stage). **b**, Pharyngeal pumping measurement over 30 seconds of wild type supplemented with or without folic acid (day 1). **c**, Brood size of WT with or without folic acid. Self-fertilizing brood size was measured during adulthood. **d**, Targeted metabolite analysis on folic acid cycle intermediates in HT115 bacteria upon folic acid and 5MTHF supplementation. No significance was observed between conditions. **e**, Mini-RNAi screen for life span of genes involved in the folic acid cycle (day 1). *dhfr-1i* and *tyms-1i* showed an increase mean lifespan >15%. **f**, Pharyngeal pumping measurement over 30 seconds in *luci* control and *dhfr-1i* treated worms(day 1). **g**, Brood size of *luci* control and *dhfr-1i* treated worms. Self-fertilizing brood size was measured during adulthood. **h**, *dhfr-1i* lifespan in the *daf-2* genetic background (from L4 stage). *dhfr-1i* has little effect on *daf-1(e1370)* lifespan. **i**, mRNA levels of selected genes implicated in methionine restriction (*dgn-1, let-23, ign-3, dapk-1* and *daf-18*) in *dhfr-1i, metr-1i* (methionine restriction control), and *luci* control backgrouns. **a,i**, n=150 per repeat per condition. **a,h**, N=3 independent biological replicates. **d**, N=2 independent biological replicates. **b,f**, n=25 worms, N=3 biological replicates. **c,g**, n=20 worms, N=3 biological replicates. **a,d**, statistics were by two-sided Mantel–Cox log-rank test. **d**, Significance was assessed using one-way anova Dunnett’s multiple comparisons test. **b,c,f,h,g**, Significance was assessed using two-side t-test. Bar shows mean± S.D, * p<0.5, ** p<0.01, ***p<0.001. (Supplementary Table 3 for statistics).

**Extended data Fig.4.**
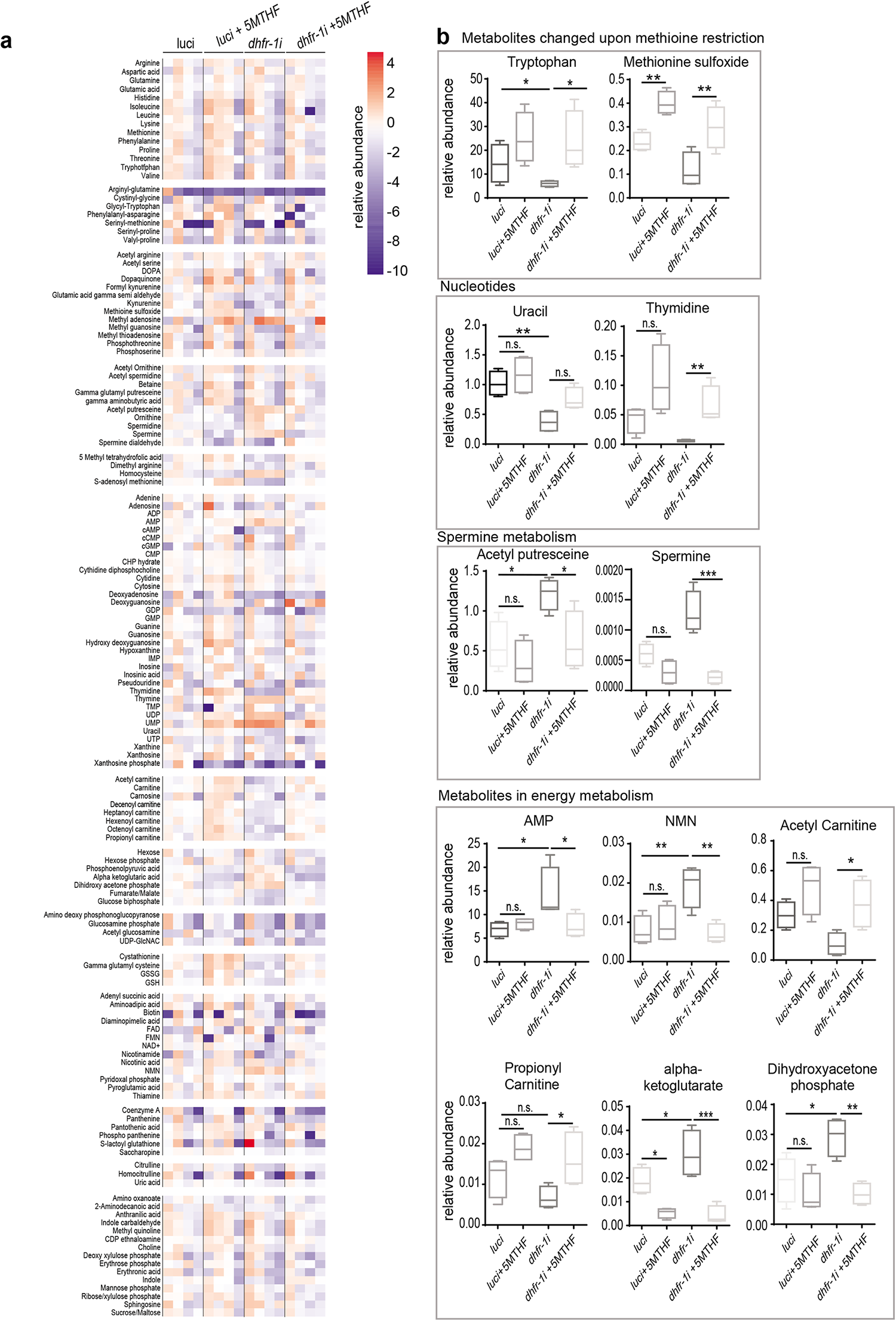
Untargeted metabolomics analysis of *dhfr-1i* with or without 5MTHF supplementation (linked to Fig 2,3). **a**, Untargeted metabolomics analysis of *luci* control *dhfr-1i* supplemented with 10 nM 5MTHF at day 1 of adulthood. Metabolites concentrations are listed in Supplementary Table 2 and manually grouped in different categories. **b**, Significant metabolites (adj p<.05) are selected from the untargeted metabolomics analysis and plotted. **a**, n=4 independent biological replicates. **a**, Metabolite concentrations are normalized to internal standard and converted to log2 for the heat map generation. **a**, statistic were determined using Fisher test and Benjamini-Hochberg correction (adj p<.05) **b**, Significance was assessed using one-way anova Dunnett’s multiple comparisons test. * p<0.5, ** p<0.01, ***p<0.001 (Supplementary Table 3 for statistics).

## Methods

Refer to Supplementary Table 3 and the Reporting Summary for all materials used to perform the experiments.

### Worm strains and culture

All strains were grown and maintained on NGM agar seeded with *E. coli* (OP50) at 20°C except for the *glp-1(e2141)ts* strain, which underwent a thermal shift to 25°C leading to germline loss. Standard procedures for culturing and maintaining strains were used^59^. Complete strain list is provided in the Supplementary Table 3.

### RNAi

Worm RNAi was conducted as described previously^60,61^. Briefly, WT wild-type worms were fed with HT115 (DE3) bacteria transformed with L4440 vector that expresses a doublestranded RNA against the targeted gene. Synchronized worms were obtained by performing an egg lay on corresponding RNAi plates containing isopropyl-β-D-thiogalactoside and ampicillin. Luciferase (L4440::Luc, i.e. *luci*) RNAi vector was used as non-targeting control. C36B1.7 (*dhfr-1*), C06A8.1(*mthf-1*), C05D11.11(*mel-32*), Y110A7A.4 (*tyms-1*), F38B6.4, K07E3.3 (*dao-3*), Y55D5A.5 (*daf-2*). R03D7.1(*metr-1*) RNAi clone were obtained from the Vidal RNAi library. The gene targeted by RNAi is indicated with an “i” after the gene name. All RNAi experiments were blinded.

### Folic acid intermediates supplementations

Folic acid and 5MTHF acid were added in aqueous solution into the NGM agar at the indicated concentrations of 10 nM. For bacteria, folic acid and 5MTHF were added to the medium and incubated 24 hours prior extraction.

### Pharyngeal pumping rate assay

Pharyngeal pumping was assessed by observing the number of pharyngeal contractions during a 30 s interval using twenty synchronized young adult worms in three biological replicates. Experiments were blinded.

### Brood size

L4 larvae (n□=□10 animals per strain) were maintained individually under standard conditions. Synchronized young adult worms were singled to 3 cm plates containing OP50. Worms were transferred to fresh plates every 24h and progeny was counted during a period of 7 days. A minimum of 10 worms were used for each genotype.

### Life span analysis

Lifespan analyses were performed at 20°C as previously reported^60,61^. Data was plotted to calculate mean, median, and maximum lifespans using Microsoft Excel 16.12 and GraphPad Prism 7 Software. For all RNAi lifespan assays 150 worms were fed with RNAi from L4 stage on. To determine significance between the lifespan curves log-rank Mantel-Cox analysis was used. Experiments were blinded.

### RNA extraction and qPCR

Synchronized day1 worms (three to four plates) were collected in Trizol (Invitrogen). Total RNA was extracted using RNeasy Mini spin column (QIAGEN). Concentration and purity of the RNA was measured by NanoDrop. cDNA was generated using iScript (Bio-Rad). qRT-PCR was performed with Power SYBR Green (Applied Biosystems) on a ViiA 7 Real-Time PCR System (Applied Biosystems). Four technical replicates were averaged for each sample per primer reaction. *cdc-42* was used as internal control. Primers are listed in Supplementary Method Table 3.

### Fluorescent microscopy

For fluorescent images of transgenic *C. elegans*, live animals were immobilized with 5 mM sodium azide and mounted on 2% agarose pads. Images were obtained with a Axio Imager Z1 Zeiss microscope. Puncta of the Q40 strain at day 7 were counted at least 30 worms in three biological replicates. Experiment was blinded.

### Thrashing assay

Motility was determined as a function of thrashing in liquid. Individual transgenic animals (Q35) at day 7 of adulthood were picked to a 10 μL drop of M9 on a microscope slide and were given a 30 s adjustment period before counting thrashing rate. Thrashes (defined as the head crossing the vertical midline of the body) were counted for 30 s. A minimal n-number of n□=□30 in three biological replicates was assayed for each genotype. Experiment was blinded.

### Metabolite extraction from worms

Synchronized young adults for each strain were collected (approx. three plates) in five biological replicates in single tubes and washed three times with buffer solution M9 and 0.1% of butyl hydroxy toluene (BHT) was added to prevent auto oxidation as previously reported on our work^18,62^. Samples were snap frozen in liquid nitrogen and stored at −80°C before use. Worm pellets were homogenized using a Qiagen tissue lyser for 30 min at 4°C. Protein concentration was determined using a BCA kit and the lysate volume corresponding to 150 μg of proteins was subjected to Bligh and Dyer extraction (chloroform: methanol, 2:1) for 1 hour at 4°C. Samples were centrifuged at maximum speed for 5 min at 4°C and supernatant was transferred into a new tube for drying. Before LC injections samples were reconstituted in 10% aqueous acetonitrile. Samples were analyzed using an untargeted method for total metabolomics and for targeted methods to evaluate the abundance of folic acid analogs and methionine cycle intermediates.

### Folic acid intermediates extraction from mouse tissues

C57BL/6 IRS1 KO and WT control were kindly provided by Linda Partridge. C57BL/6 *Irs1^-/-^* KO mice were originally obtained from Prof. Dominic Withers’ lab (Imperial College, London) and were bred on-site at the mouse facility in Max Planck Institute for Biology of Ageing, Cologne. Mouse experiments were performed according to the guidelines and approval of LANUV [Landesamt für Natur, Umwelt und Verbraucherschutz Nordrhein-Westfalen (State Agency for Nature, Environment and Consumer Protection North Rhine-Westphalia), VSG 84-02.04.2013.A158]. The animals were sacrificed at the age of 24 months and tissues were snap-frozen in liquid nitrogen and kept at −80°C. Tissues underwent similar extraction as described for the worms. Tissues were homogenized using a Qiagen tissue lyser for 30 min at 4°C. Protein concentration was determined using a BCA kit and the lysate volume corresponding to 150 μg of protein was subjected to Bligh and Dyer extraction (chloroform: methanol, 2:1) for 1 hour at 4°C. Samples were centrifuged at maximum speed for 5 min at 4°C and supernatant was transferred into a new tube for drying. Before LC injections samples were reconstituted in 10% aqueous acetonitrile. Samples were analyzed using a targeted method to assess the abundance of folic acid and methionine cycle intermediates.

### Untargeted metabolomics

Analytes were separated using an UHPLC system (Vanquish, Thermo Fisher Scientific, Bremen, Germany) coupled to an HRAM mass spectrometer (Q-Exactive Plus, Thermo Fischer Scientific GmbH, Bremen, Germany) using a modified RP-MS method from Wang L *et al*^63^. Briefly, two microliters of the sample extract were injected into a X Select HSS T3 XP column, 100 Å, 2.5 μm, 2.1 mm x 100 mm (Waters), using a binary system A water with 0.1% formic acid, B: acetonitrile with 0.1 formic acid with a flowrate of 0.1 mL/min, with the column temperature kept at 30 °C. Gradient elution was conducted as follows: isocratic step at 0.1 % eluent B for 0.3 min, gradient increase up to 2% eluent B in 2 min, then increase up to 30% eluent B in 6 min and to 95% eluent B in 7 min, isocratic step at 95% eluent B for 2 min. Gradient decreases to 0.1 % eluent B in 3 min and held at 0.1% eluent B for 5 min. Mass spectra were recorded from 100-800 *m/z* at a mass resolution of 70,000 at *m/z* 400 in both positive and negative ion modes using data dependent acquisition (Top 3, dynamic exclusion list 10 seconds). Tandem mass spectra were acquired by performing CID (isolation 1,5 a.u., stepped collision energy 20 and 80 NCE). The *m/z* of Leucine enkephaline was used as lock mass. Sample injection order was randomized to minimize the effect of instrumental signal drift. MS data analysis was performed using Xcalibur software 4.0.

### Compound identification and quantification

Metabolite search was performed using Compound discoverer 2.0 and *m/z C*loud as online databases, considering precursor ions with a deviation > 5 ppm, 0.3 min maximum retention time shift, minimum peak intensity 100000, intensity tolerance 10, FT fragment mass tolerance 0.0025 Da, group covariance [%] less than 30, *p*-value less than 0.05 and area Max greater or equal to 10000. Metabolites are correctly identified when at least two specific fragments are found in the MS^2^ spectra. Because of the high mass accuracy >3 ppm, predicted elemental compositions of the unknown features were submitted to other online databases such as Chemspider (http://www.chemspider.com/), HMDB (http://www.hmdb.org/), KEGG (http://www.genome.jp/kegg/), METLIN (http://metlin.scripps.edu/). Unassigned features were additionally submitted to PIUMet algorithm for pathway prediction (http://fraenkel-nsf.csbi.mit.edu/piumet2/). The output was processed using R packages ‘gplot’ in order to visualize the cluster of metabolites and to highlight the connection between the predicted proteins and enzymes.

Quantification was performed using Trace finder 4.1, using genesis detection algorithm, nearest RT, S/N threshold 8, min peak height (S/N) equal to 3, peak S/N cutoff 2.00, valley rise 2%, valley S/N 1.10. Relative quantification was obtained by dividing the area of individual metabolites to spiked internal standards (Leucine enkephaline, myrystic acid and cysteamine sodium salt).

### Targeted analysis of folic acid intermediates

Identification and relative quantification of folic acid intermediates were performed on a triple quadrupole mass spectrometer (QqQMS) (TSQ Altis, ThermoFisher Scientific GmbH, Bremen, Germany), as previously published by our group^18^. Data was analyzed using Xcalibur version 4.0. Quantification was performed using Trace finder 4.1, using genesis detection algorithm, nearest RT, S/N threshold 8, min peak height (S/N) equal to 3, peak S/N cutoff 2.00, valley rise 2%, valley S/N 1.10. The relative response for each folate species was calculated by dividing the peak area of the analyte to the internal standard peak area (pteridinic acid) and further normalized to protein concentration.

### Targeted analysis of methionine cycle intermediates

Methionine intermediates were identified and quantified using a high resolution accurate mass (HRAM) mass spectrometer (Q-Exactive Plus, Thermo Fischer Scientic GmbH, Bremen,Germany) coupled with an UHPLC system (Vanquish, Thermo Fisher Scientific, Bremen, Germany). Analytes were separated using a X Select HSS T3 XP column, 100 Å, 2.5 μm, 2.1 mm x 100 mm (Waters), using a binary system A water with 0.1% formic acid, B: acetonitrile with 0.1 formic acid with a flowrate of 0.1 mL/min and the column temperature was kept at 30°C. Gradient elution was conducted as for untargeted metabolomics analysis. Methionine cycle intermediates were identified using a Targeted-SIM (t-SIM) with a resolution of 70,000, 5e^4^ AGC target, 200 ms injection time and 1.0 *m/z* isolation window. The following ions were quantified: Methionine-> 149.0.5084, s-adenosyl methionine −> 398.13724, Homocysteine −>135.03540, S adenosyl homocysteine −>385.12800. Quantification was performed using Trace finder 4.1, using genesis detection algorithm, nearest RT, S/N threshold 8, min peak height (S/N) equal to 3, peak S/N cutoff 2.00, valley rise 2%, valley S/N 1.10. The relative response for each methionine intermediate was calculated by dividing the peak area of the analyte to the internal standard peak area and further normalized to protein concentration.

### 5MTHF incorporation

In order to elucidate the *in vivo* kinetics of 5MTHF in a dynamic metabolite flux analysis a single pulse of isotopic labelled 5 MTHF/Glutamic acid 13C15N in a concentration of 5 μM was added to synchronized day 1 young adult worms on NGM plates. RNAi treatment for *dhfr-1* knockdown was performed prior to the pulse. Worms were collected for metabolite extraction at 7 time points (0, 10, 30, 60, 120, 240, 360 minutes). Metabolites extraction was conducted as described above. The transitions of 13C15N label 5MTHF were added to the initial method.

### Statistical data

GraphPad Prism Version 7.0c software was used for graphics and statistical testing. Metabolomics data sets were analyzed using Fisher test and Hochberg-Benjamin false discovery rate test. Individual metabolites and targeted metabolomics were analyzed using one-way-anova and Dunnett’s correction test. Lifespan experiments were analyzed using log-rank Mantel-Cox test.

### Data availability

The data that support the findings of this study are available within the paper (and supplementary information files) or from the corresponding author upon reasonable request.

