## Supplementary Table 3 for "Regulation of the one carbon folate cycle as a shared metabolic signature of longevity": Supplementary Table 3.docx

Regulation of the one carbon folate cycle as a shared metabolic signature of longevity

**Authors:** 1Andrea Annibal, 1Rebecca George Tharyan ,1Maribel Schoenewolff, 1Hannah Tam, 1Christian Latza, Markus Max Karl Auler 1,2,3Adam Antebi

1Max Planck Institute for Biology of Ageing, Joseph Stelzmann Str. 9b, 50931, Cologne, Germany

2Cologne Excellence Cluster on Cellular Stress Responses in Aging-Associated Diseases (CECAD), University of Cologne, 50931, Cologne, Germany

3Corresponding author

**Table1: Strains used in this study**

| **AA numbers/Strains:** | **Genotype:** |
| --- | --- |
|  | ***N2 (wild type,* WT*)*** |
| **AA3561** | *glp-1 (e2141)ts III* |
| **CB1370** | daf-2(e1370) III |
| **MQ887** | isp-1(qm150) IV |
| **DA465** | eat-2 (ad465) II |
| **AA60** | daf-16(mgDf50) I |
| **AA4776** | N2; rmIs132[P(unc-54) Q35::YFP] |
| **AM141** | rmIs133[P(unc-54) Q40::YFP] |

**Table 2: Lifespan Analysis**

| **Figure 2a** | | | | | | | | | | |
| --- | --- | --- | --- | --- | --- | --- | --- | --- | --- | --- |
| **Strains** | **Treatment/RNAi** | Mean LS | Mean Diff. | **Max LS** | **Max Diff.** | Median LS | **Median Diff.** | **Log-rank(Mantel-Cox) test** | **Comparison** | **Replicates** |
| WT | *luciferase* | 18.54 | 0.00% | 28 | 0.00% | 17 | 0.00% |  |  | BR1 |
| WT | *dhfr-1i* | 23.87 | 28.77% | 36 | 28.57% | 24 | 24.18% | < 0,0001 | vs *luciferase* |  |
| WT | *tyms-1i* | 22.28 | 20.18% | 36 | 28.57% | 21 | 23.53% | 0.0001 | vs *luciferase* |  |
| WT | *daf-2i* | 32.58 | 75.75% | 58 | 107.14% | 32 | 88.24% | < 0,0001 | vs *luciferase* |  |
| WT | *luciferase* | 16.42 | 0.00% | 30 | 0.00% | 17 | 0.00% |  |  | BR2 |
| WT | *dhfr-1i* | 21.57 | 31.41% | 36 | 20.00% | 21 | 23.53% | 0.0002 | vs *luciferase* |  |
| WT | *tyms-1i* | 21.28 | 29.65% | 34 | 13.33% | 21 | 23.53% | 0.0003 | vs *luciferase* |  |
| WT | *daf-2i* | 30.18 | 83.83% | 60 | 100.00% | 28 | 64.71% | < 0,0001 | vs *luciferase* |  |
| WT | *luciferase* | 16.00 | 0.00% | 28 | 0.00% | 17 | 0.00% |  |  | BR3 |
| WT | *dhfr-1i* | 21.36 | 30.10% | 34 | 21.43% | 21 | 23.53% | 0.0002 | vs *luciferase* |  |
| WT | *tyms-1i* | 20.88 | 27.18% | 36 | 28.57% | 21 | 23.53% | 0.0007 | vs *luciferase* |  |
| WT | *daf-2i* | 28.74 | 75.07% | 56 | 100.00% | 28 | 64.71% | < 0,0001 | vs *luciferase* |  |
| **Figure 2b, Extended data Fig. 3a** | | | | | | | | | | |
| **Strains** | **Treatment/RNAi** | Mean LS | Mean Diff. | **Max LS** | **Max Diff.** | Median LS | **Median Diff.** | **Log-rank(Mantel-Cox) test** | **Comparison** | **Replicates** |
| WT | *luciferase* | 21.01 | 0.00% | 29.00 | 0.00% | 23.00 | 0.00% |  |  | BR1 |
| WT | *luciferase* +10 nM FA | 19.89 | -5.34% | 29.00 | 0.00% | 21.00 | -8.70% | 0.0621 | vs *luciferase* |  |
| WT | *luciferase* +10 nM 5MTHF | 21.01 | 0.02% | 30.00 | 3.45% | 21.00 | -8.70% | 0.9067 | vs *luciferase* |  |
| WT | *dhfr-1i* | 25.79 | 22.75% | 39.00 | 34.48% | 25.00 | 8.70% | <0.0001 | vs *luciferase* |  |
| WT | *dhfr-1i* + 10nM 5MTHF | 20.57 | -2.10% | 29.00 | 0.00% | 21.00 | -8.70% | <0.0001 | vs *dhfr-1i* |  |
| WT | *luciferase* | 20.30 | 0.00% | 29.00 | 0.00% | 19.00 | 0.00% |  |  | BR2 |
| WT | *luciferase* +10 nM FA | 19.20 | -5.40% | 29.00 | 0.00% | 19.00 | 0.00% | 0.238 | vs *luciferase* |  |
| WT | *luciferase* +10 nM 5MTHF | 20.17 | -0.62% | 30.00 | 3.45% | 19.00 | 0.00% | 0.5604 | vs *luciferase* |  |
| WT | *dhfr-1i* | 23.82 | 17.36% | 39.00 | 34.48% | 23.00 | 21.05% | <0.0001 | vs *luciferase* |  |
| WT | *dhfr-1i* + 10nM 5MTHF | 21.19 | 4.39% | 29.00 | 0.00% | 21.00 | 10.53% | 0.0017 | vs *dhfr-1i* |  |
| WT | *luciferase* | 22.02 | 0.00% | 30.00 | 0.00% | 23.00 | 0.00% |  |  | BR3 |
| WT | *luciferase* +10 nM FA | 22.17 | 0.68% | 32.00 | 0.00% | 21.00 | 0.00% | 0.6638 | vs *luciferase* |  |
| WT | *luciferase* +10 nM 5MTHF | 21.95 | -0.30% | 32.00 | 0.00% | 21.00 | 0.00% | 0.8677 | Vs *luciferase* |  |
| WT | *dhfr-1i* | 25.78 | 22.80% | 39.00 | 21.88% | 25.00 | 8.69% | 0.0146 | vs *luciferase* |  |
| WT | *dhfr-1i* + 10nM 5MTHF | 20.80 | -5.53% | 32.00 | 0.00% | 21.00 | -8.70% | 0.0029 | vs *dhfr-1i* |  |
| **Figure 4a,b** | | | | | | | | | | |
| **Strains** | **Treatment/RNAi** | Mean LS | Mean Diff. | **Max LS** | **Max Diff.** | Median LS | **Median Diff.** | **Log-rank(Mantel-Cox) test** | **Comparison** | **Replicates** |
| WT |  | 21.90 | 0.00% | 31.00 | 0.00% | 22.00 | 0.00% |  |  | BR1 |
| *daf-2(e1370)* |  | 41.93 | 91.55% | 62.00 | 100.00% | 41.00 | 86.36% | <0.0001 | vs WT |  |
| *daf-2(e1370)* | +10nM 5MTHF | 38.94 | 77.91% | 57.00 | 83.87% | 22.00 | 77.27% | 0.0208 | vs *daf-2(e1370)* |  |
| *isp-1(qm150)* |  | 29.86 | 36.39% | 44.00 | 41.94% | 29.00 | 31.82% | <0.0001 | vs WT |  |
| *isp-1(qm150)* | +10nM 5MTHF | 27.01 | 23.40% | 34.00 | 9.68% | 29.00 | 31.82% | 0.001 | vs *isp-1(qm150)* |  |
| WT |  | 23.05 | 0.00% | 31.00 | 0.00% | 24.00 | 0.00% |  |  | BR2 |
| *daf-2(e1370)* |  | 42.87 | 86.00% | 64.00 | 106.45% | 41.00 | 70.83% | <0.0001 | vs WT |  |
| *daf-2(e1370)* | +10nM 5MTHF | 39.21 | 70.11% | 57.00 | 83.87% | 37.00 | 54.17% | 0.013 | vs *daf-2(e1370)* |  |
| *isp-1(qm150)* |  | 30.15 | 30.78% | 44.00 | 41.94% | 31.00 | 29.17% | <0.0001 | vs WT |  |
| *isp-1(qm150)* | +10nM 5MTHF | 27.19 | 17.97% | 34.00 | 9.68% | 29.00 | 20.83% | 0.0011 | vs *isp-1(qm150)* |  |
| WT |  | 20.68 | 0.00% | 31.00 | 0.00% | 24.00 | 0.00% |  |  | BR3 |
| *daf-2(e1370)* |  | 41.22 | 99.31% | 62.00 | 100.00% | 41.00 | 70.83% | <0.0001 | vs WT |  |
| *daf-2(e1370)* | *+10nM 5MTHF* | 29.21 | 41.27% | 48.00 | 54.84% | 35.00 | 45.83% | <0.0001 | vs *daf-2(e1370)* |  |
| *isp-1(qm150)* |  | 26.59 | 28.59% | 44.00 | 41.94% | 26.00 | 8.33% | <0.0001 | vs WT |  |
| *isp-1(qm150)* | *+10nM 5MTHF* | 23.51 | 13.67% | 34.00 | 9.68% | 24.00 | 0.00% | 0.0019 | vs *isp-1(qm150)* |  |
| **Figure 4e** | | | | | | | | | | |
| **Strains** | **Treatment/RNAi** | Mean LS | Mean Diff. | **Max LS** | **Max Diff.** | Median LS | **Median Diff.** | **Log-rank(Mantel-Cox) test** | **Comparison** | **Replicates** |
| WT | *luciferase* | 21.67 | 0.00% | 32.00 | 0.00% | 21.00 | 0.00% |  |  | BR1 |
| WT | *dhfr-1i* | 25.36 | 17.03% | 39.00 | 21.88% | 28.00 | 33.33% | <0.0001 | vs *luciferase* |  |
| *daf-16(mgDf50)* | *luciferase* | 10.48 | -51.62% | 21.00 | -34.38% | 9.00 | -57.14% | <0.0001 | vs *luciferase* |  |
| *daf-16(mgDf50)* | *luciferase + 10nM 5MTHF* | 9.91 | -54.27% | 23.00 | -28.13% | 9.00 | -57.14% | 0.5068 | vs *daf-16(mgDf50) luciferase* |  |
| *daf-16(mgDf50)* | *dhfr-1i* | 12.26 | -43.40% | 25.00 | -21.88% | 14.00 | -33.33% | <0.0001 | vs *daf-16(mgDf50) luciferase* |  |
| *daf-16(mgDf50)* | *dhfr-1i + 10nM 5MTHF* | 10.77 | -50.27% | 23.00 | -28.13% | 11.00 | -47.62% | 0.0005 | vs *daf-16(mgDf50) dhfr-1i* |  |
| WT | *luciferase* | 21.21 | 0.00% | 32.00 | 0.00% | 21.00 | 0.00% |  |  | BR2 |
| WT | *dhfr-1i* | 23.51 | 10.84% | 39.00 | 21.88% | 23.00 | 9.52% | <0.0001 | vs *luciferase* |  |
| *daf-16(mgDf50)* | *luciferase* | 11.33 | -46.59% | 23.00 | -28.13% | 11.00 | -47.62% | <0.0001 | vs *luciferase* |  |
| *daf-16(mgDf50)* | *Luciferase + 10nM 5MTHF* | 12.49 | -41.12% | 23.00 | -28.13% | 14.00 | -33.33% | 0.1181 | vs *daf-16(mgDf50) luciferase* |  |
| *daf-16(mgDf50)* | *dhfr-1i* | 12.16 | -42.69% | 25.00 | -21.88% | 11.00 | -47.62% | 0.3298 | vs *daf-16(mgDf50) luciferase* |  |
| *daf-16(mgDf50)* | *dhfr-1i + 10nM 5MTHF* | 9.77 | -53.91% | 23.00 | -28.13% | 9.00 | -57.14% | 0.9341 | vs *daf-16(mgDf50) dhfr-1i* |  |
| WT | *luciferase* | 18.94 | 0.00% | 30.00 | 0.00% | 19.00 | 0.00% |  |  | BR3 |
| WT | *dhfr-1i* | 22.61 | 19.62% | 35.00 | 16.67% | 23.00 | 21.05% | 0.0029 | vs *luciferase* |  |
| *daf-16(mgDf50)* | *luciferase* | 11.35 | -39.96% | 23.00 | -23.33% | 11.00 | -42.11% | <0.0001 | vs *luciferase* |  |
| *daf-16(mgDf50)* | *luciferase + 10nM 5MTHF* | 10.12 | -46.43% | 23.00 | -23.33% | 9.00 | -52.63% | 0.1209 | vs *daf-16(mgDf50) luciferase* |  |
| *daf-16(mgDf50)* | *dhfr-1i* | 12.53 | -33.69% | 25.00 | -16.67% | 11.00 | -42.11% | 0.0978 | vs *daf-16(mgDf50) luciferase* |  |
| *daf-16(mgDf50)* | *dhfr-1i + 10nM 5MTHF* | 10.72 | -43.27% | 23.00 | -23.33% | 9.00 | -52.63% | 0.0019 | vs *daf-16(mgDf50) dhfr-1i* |  |
| **Extended data Fig 3e** | | | | | | | | | | |
| **Strains** | **Treatment/RNAi** | **Mean LS** | **Mean Diff.** | **Max LS** | **Max Diff.** | **Median LS** | **Median Diff.** | **Log-rank(Mantel-Cox) test** | **Comparison** | **Replicate** |
| WT | *luciferase* | 18.54 | 0.00% | 28 | 0.00% | 17 | 0.00% |  |  | BR1 |
| WT | *mthfr-1i* | 20.61 | 3.28% | 30 | 0.00% | 21 | 0.00% | 0.075 | vs *luciferase* |  |
| WT | *mel-32i* | 17.72 | -0.11% | 28 | 0.00% | 17 | 0.00% | 0.356 | vs *luciferase* |  |
| WT | *tyms-1i* | 24.12 | 28.60% | 36 | 21.43% | 24 | 23.53% | < 0,0001 | vs *luciferase* |  |
| WT | *dhfr-1i* | 23.89 | 36.04% | 36 | 28.57% | 24 | 23.53% | < 0,0001 | vs *luciferase* |  |
| WT | *F38B6.4i* | 17.81 | 12.89% | 28 | 7.14% | 19 | 0.00% | 0.3076 | vs *luciferase* |  |
| WT | *daf-2i* | 33.68 | 100.20% | 58 | 114.29% | 32 | 88.24% | < 0,0001 | vs *luciferase* |  |
| WT | *dao-3i* | 16.05 | 9.18% | 30 | 0.00% | 15 | 11.76% | 0.0718 | vs *luciferase* |  |
| WT | *luciferase* | 16.98 | 0.00% | 30 | 0.00% | 17 | 0.00% |  |  | BR2 |
| WT | *mthfr-1i* | 17.08 | 0.60% | 28 | -6.67% | 17 | 0.00% | 0.8436 | vs *luciferase* |  |
| WT | *mel-32i* | 16.52 | -2.70% | 28 | -6.67% | 17 | 0.00% | 0.5554 | vs *luciferase* |  |
| WT | *tyms-1i* | 21.27 | 25.26% | 34 | 13.33% | 21 | 23.53% | 0.0001 | vs *luciferase* |  |
| WT | *dhfr-1i* | 22.50 | 32.51% | 36 | 20.00% | 21 | 23.53% | < 0,0001 | vs *luciferase* |  |
| WT | *F38B6.4i* | 18.67 | 9.96% | 30 | 0.00% | 17 | 0.00% | 0.129 | vs *luciferase* |  |
| WT | *daf-2i* | 33.11 | 95.02% | 60 | 100.00% | 32 | 88.24% | < 0,0001 | vs *luciferase* |  |
| WT | *dao-3i* | 18.06 | 6.35% | 28 | -6.67% | 19 | 11.76% | 0.583 | vs *luciferase* |  |
| **Extended data Fig. 3h** | | | | | | | | | | |
| **Strains** | **Treatment/RNAi** | **Mean LS** | **Mean Diff.** | **Max LS** | **Max Diff.** | **Median LS** | **Median Diff.** | **Log-rank(Mantel-Cox) test** | **Comparison** | **Replicate** |
| WT | *luciferase* | 16.25 | 0% | 30 | 0% | 19 | 0% |  |  | BR1 |
| WT | *dhfr-1i* | 21.63 | 33.14% | 36 | 20% | 22 | 15.79% | 0.0004 | vs *luciferase* |  |
| *daf-2(e1370)* | *luciferase* | 30.35 | 86.79% | 54 | 80% | 32 | 68.42% | <0.0001 | vs *luciferase* |  |
| *daf-2(e1370)* | *dhfr-1i* | 29.56 | 81.97% | 54 | 80% | 30 | 57.89% | <0.0001 | vs *luciferase* |  |
|  |  |  |  |  |  |  |  | 0.2610 | vs *daf-2(e1370)* |  |
| WT | *luciferase* | 18.53 | 0% | 28 | 0% | 17 | 0% |  |  | BR2 |
| WT | *dhfr-1i* | 23.91 | 31.20% | 35 | 25% | 22 | 29.41% | 0.0004 | vs *luciferase* |  |
| *daf-2(e1370)* | *luciferase* | 31.43 | 69.59% | 58 | 107.14% | 30 | 76.47% | <0.0001 | vs *luciferase* |  |
| *daf-2(e1370)* | *dhfr-1i* | 29.97 | 61.67% | 58 | 107.14% | 30 | 76.47% | <0.0001 | vs *luciferase* |  |
|  |  |  |  |  |  |  |  | 0.3458 | vs *daf-2(e1370)* |  |

| **Table 3: qPCR primers** | |  |
| --- | --- | --- |
| **Gene name** | **Forward primer sequence** | **Reverse primer sequence** |
| ***dhfr-1*** | TTTCGGAGCCACCATTTGTG | ACCAAGTTTTCTCGCAACGC |
| ***mel-32*** | TTGTTGACTTGCGCCCAATC | TGGGCACGTATTCTTGTTGC |
| ***mthfr-1*** | ATGGGGAAACAGTTCATCGC | TTGATGAACACGCGCTTGAC |
| ***dao-3*** | ATCAATGCTGGACGTTTGGC | TTCGAGCGTCCCAAAACAAC |
| ***F38B6.4*** | AGGTTTCTGCGTTGGCTTTC | AACGTTTGTGGTCCTTTCCG |
| ***dapk-1*** | TGGCAACATTTGTGCACGAG | ACTGTATGCTGAGCAAAGCG |
| ***ing-3*** | TACTGATGACACCGACGTTCTC | ACACCACAAACGCCGAATTC |
| ***daf-18*** | AAATACGGCCGGTAACATGC | TGGTCAACGAAGGCTTTTGC |
| ***let-23*** | CGCTGAAATGGTTGACATGC | TTTGGTTTGCAGCTCGGATG |
| ***dgn-1*** | AAATCAAGAGGACGCAAGCC | AGTGTTGTTGGCTCCACATG |

**Table 4: Features submitted to PIUMET (linked to Fig. 1c)**

| Molecular weight | Ion mode | Retention time |
| --- | --- | --- |
| 304.2038 | positive | 13.279 |
| 515.3015 | positive | 16.277 |
| 547.3632 | positive | 21.297 |
| 545.3481 | positive | 18.407 |
| 571.3638 | positive | 12.865 |
| 473.2481 | positive | 9.193 |
| 505.3173 | positive | 18.214 |
| 503.3008 | positive | 17.255 |
| 501.2798 | positive | 9.157 |
| 499.2689 | positive | 20.962 |
| 529.3117 | positive | 12.401 |
| 302.1882 | positive | 13.22 |
| 451.2699 | positive | 21.071 |
| 479.3007 | positive | 16.53 |
| 477.2855 | positive | 7.941 |
| 691.5157 | positive | 20.782 |
| 733.5593 | positive | 20.893 |
| 783.5728 | positive | 21.425 |
| 777.5297 | positive | 19.306 |
| 430.3083 | positive | 12.689 |
| 761.5826 | positive | 20.933 |
| 689.498 | positive | 17.538 |
| 687.4834 | positive | 20.952 |
| 737.4953 | positive | 20.323 |
| 705.5304 | positive | 18.118 |
| 767.5449 | positive | 21.292 |
| 797.5915 | positive | 20.94 |
| 795.5716 | positive | 21.057 |
| 713.4955 | positive | 20.185 |
| 787.512 | positive | 20.737 |
| 120.0299 | positive | 2.14 |
| 727.55 | positive | 21.141 |
| 288.1725 | positive | 2.609 |
| 300.2089 | positive | 11.492 |
| 584.4441 | positive | 12.035 |
| 552.4748 | positive | 10.553 |
| 752.6327 | positive | 21.721 |
| 519.3207 | positive | 17.227 |
| 517.3163 | positive | 16.946 |
| 132.0774 | negative | 12.903 |
| 132.0775 | negative | 13.467 |
| 475.2696 | negative | 16.709 |
| 529.3168 | negative | 13.26 |
| 481.3168 | negative | 21.589 |
| 477.2855 | negative | 17.28 |
| 221.1048 | negative | 13.533 |
| 219.1103 | negative | 9.062 |
| 731.5445 | negative | 20.746 |
| 745.5598 | negative | 19.505 |
| 717.5306 | negative | 21.273 |
| 731.5793 | negative | 20.77 |
| 316.2191 | negative | 12.071 |
| 230.0192 | negative | 13.141 |
| 336.074 | negative | 8.142 |
| 536.0445 | negative | 2.428 |
| 147.0532 | negative | 3.493 |
| 515.3104 | negative | 13.023 |
| 543.3418 | negative | 13.435 |
| 541.3285 | negative | 17.104 |
| 567.3442 | negative | 17.645 |

**Table 5: Targeted metabolomics analysis of folates in longevity mutants**

| **Figure 1d** | | | | | |
| --- | --- | --- | --- | --- | --- |
|  | **Average peak area normalized to control (WT)** | | | | |
| **Compounds** | **WT** | ***daf-2(e1370)*** | ***isp-1(qm150)*** | ***eat-2(ad465)*** | ***glp-1(e2141ts)*** |
| FA | 1.0000 | 3.9657 | 1.8359 | 2.2170 | 0.9646 |
| THF | 1.0000 | 5.2470 | 1.8451 | 1.9509 | 1.4456 |
| 5MTHF | 1.0000 | 0.4155 | 0.5234 | 0.3232 | 1.1651 |
| MN | 1.0000 | 1.4457 | 3.5562 | 3.2902 | 1.2650 |
| ML | 1.0000 | 0.1228 | 0.7649 | 0.5316 | 1.2557 |
| FO | 1.0000 | 0.6266 | 1.3589 | 0.4206 | 0.8029 |
| **Compounds** |  | ***daf-2(e1370)***  ***vs WT*** | ***isp-1(qm150)***  ***vs WT*** | ***eat-2(ad465)***  ***vs WT*** | ***glp-1(e2141ts)***  ***vs WT*** |
| FA |  | 0.0017** | 0.003** | 0.0029** | 0.9060 |
| THF |  | 0.0002*** | 0.019* | 0.0254* | 0.0211* |
| 5MTHF |  | 0.0002*** | 0.0291* | 0.0002*** | 0.2447 |
| ML |  | 0.0303* | 0.0033** | <0.0001*** | 0.2571 |
| MN |  | <0.0001*** | 0.0987 | 0.0032** | 0.3388 |
| FO |  | 0.0012** | 0.1126 | 0.0274* | 0.3756 |
| **Figure 2c** | | | | | |
|  | **Average peak area normalized to control (WT)** | | | | |
| **Compounds** | **Luciferase** | **Luciferase +10nM FA** | **Luciferase +10nM 5MTHF** | ***dhfr-1i*** | ***dhfr-1i* +10nM 5MTHF** |
| FA | 1.0000 | 0.9621 | 1.0330 | 2.0318 | 1.8614 |
| THF | 1.0000 | 1.1318 | 2.1130 | <LOD | 1.1314 |
| 5MTHF | 1.0000 | 0.7106 | 1.5118 | 0.4364 | 0.9588 |
| MN | 1.0000 | 0.8732 | 1.4748 | 0.3445 | 0.9179 |
| ML | 1.0000 | 1.0180 | 1.0430 | 0.5353 | 0.9549 |
| FO | 1.0000 | 0.7715 | 0.8714 | 0.4964 | 0.7587 |
| **Compounds** |  | **Luciferase vs +10nM FA** | **Luciferase vs +10nM 5MTHF** | **Luciferase vs *dhfr-1i*** | ***dhfr-1i* vs *dhfr-1i* +10nM 5MTHF** |
| FA |  | 0.0001 | 0.0001 | 0.0001 | 0.6577 |
| THF |  | 0.7449 | 0.0001 | N.D | N.D |
| 5MTHF |  | 0.0405 | 0.0004 | 0.0001 | 0.0003 |
| MN |  | 0.3897 | 0.0001 | 0.0001 | 0.0001 |
| ML |  | 0.9997 | 0.9925 | 0.0015 | 0.0038 |
| FO |  | 0.1707 | 0.6493 | 0.0009 | 0.0965 |
|  |  |  |  |  | N.D: refers to not detected |
| **Figure 3b,c** |  |  |  |  |  |
|  | **Average peak area normalized to control (WT, Luciferase)** | | | | |
| **Compounds** | **Luciferase** | **Luciferase + 10nM FA** | **Luciferase + 10nM 5MTHF** | **Luciferase vs *dhfr-1i*** | ***dhfr-1i* vs *dhfr-1i* +10nM 5MTHF** |
| Methionine | 1.0000 | 0.8567 | 1.4985 | 0.5713 | 0.9839 |
| S-adenosyl methionine | 1.0000 | 0.7471 | 1.4166 | 0.5764 | 1.1430 |
| S adenosyl Homocysteine | 1.0000 | 1.4537 | 0.7469 | 1.5948 | 0.6804 |
| Homocysteine | 1.0000 | 1.4126 | 0.6632 | 1.6391 | 0.4557 |
| **Compounds** |  | **Luciferase vs +10nM FA** | **Luciferase vs +10nM 5MTHF** | **Luciferase vs *dhfr-1i*** | ***dhfr-1i* vs *dhfr-1i* +10nM 5MTHF** |
| Methionine |  | 0.2930 | 0.0001 | 0.0005 | 0.0007 |
| S-adenosyl methionine |  | 0.0084 | 0.0001 | 0.0001 | 0.0001 |
| S adenosyl Homocysteine |  | 0.0008 | 0.0517 | 0.0001 | 0.0001 |
| Homocysteine |  | 0.0001 | 0.0007 | 0.0001 | 0.0001 |
